# Combining Focused Identification of Germplasm and Core Collection Strategies to Identify Genebank Accessions for Central European Soybean Breeding

**DOI:** 10.1101/848978

**Authors:** Max Haupt, Karl Schmid

**Affiliations:** Institute of Plant Breeding, Seed Science and Population Genetics, University of Hohenheim, Stuttgart, Germany

**Keywords:** Soybean, adaptation, core collection, plant breeding, genetic diversity, genebank

## Abstract

Environmental adaptation of crops is essential for reliable agricultural production and an important breeding objectives. Genbanks provide genetic variation for the improvement of modern varieties, but the selection of suitable germplasm is frequently impeded by incomplete phenotypic data. We address this bottleneck by combining a *Focused Identification of Germplasm Strategy* (FIGS) with core collection methodology to select soybean (*Glycine max*) germplasm for Central European breeding from a collection of >17,000 accessions. By focussing on environmental adaptation to high-latitude cold regions, we selected an ‘environmental precore’ of 3,663 accessions using environmental data and compared the Donor Population of Environments (DPE) in Asia and the Target Population of Environments (TPE) in Central Europe in the present and in 2070. Using SNP genotypes we reduced the precore into two diverse core collections of 183 and 366 of accessions as diversity panels for evaluation in high-latitude cold regions. Tests of genetic differentiation between precore and core collections revealed differentiation signatures in genomic regions that control maturity, and novel candidate loci for environmental adaptation demonstrating the potential of diversity panels for studying environmental adaptation. Objective-driven core collections increase germplasm utilization for abiotic adaptation by breeding for a rapidly changing climate, or *de novo* adaptation of crop species to expand cultivation ranges.

## INTRODUCTION

Many crop species are cultivated outside their center of domestication (Diamond 2002). An expansion of the cultivation range requires adaptation to local biotic and abiotic environmental conditions, which can be limited by reduced genetic variation as a result of variation ‘left behind’ in the center of origin due to founder effects (Tanksley and McCouch 1997; Hyten et al. 2006). Nevertheless, numerous crops were successfully adapted to novel environments by natural and human selection that contributed to the rapid expansion of farming beyond centers of domestication (Harlan 1992). In current plant breeding programs, reduced levels of genetic variation in elite breeding populations limit breeding progress and the adaptation to rapidly changing environmental conditions. Plant genetic resources stored in large *ex situ* germplasm collections are an important source of novel, potentially useful genetic variation to achieve these goals (Wang et al. 2017a).

The efficient utilization of plant genetic resources by breeders and researchers is frequently limited by incomplete phenotypic data for traits of interest, which makes the targeted selection of putative useful accessions from large germplasm collections very challenging. The *Focused Identification of Germplasm Strategy* (FIGS) promises a solution to this phenotyping bottleneck for traits associated with environmental adaptation. FIGS is based on the premise that native landraces evolved during centuries of cultivation in response to local eco-geographic conditions which led to local adaptation to biotic and abiotic conditions. Genomic footprints of this adaptation should be detectable if environmental parameters describing plant germplasm collection sites are used as selection criteria to ‘focus in’ on a subset of accessions that potentially harbor phenotypic variation in a target trait. The comprehensive phenotypic and genomic evaluation of focused subsets is more resource efficient than of large collections (Street et al. 2008; Sanders et al. 2013). FIGS has been successfully used in a number of crops including *Hordeum vulgare* for agro-morphological characteristics and time of heading and maturity (Endresen 2010), *Triticum aestivum* for resistances to powdery mildew (Bhullar et al. 2009) and stem rust (Bari et al. 2012), and *Vicia faba* for drought tolerance (Khazaei et al. 2013). A second tool in the characterization of germplasm collections are core collections. They are small subsets of larger collections that maximize genetic and phenotypic diversity (Frankel 1984). Similar to trait-focused subsets, core collections provide a resource efficient means for further evaluation and utilization and have been established for many crops based on passport information, agronomic and morphological data as well as genetic diversity to serve varying objectives (Odong et al. 2013).

The soybean (*Glycine max*) is a typical example of a crop whose cultivation expanded outside its center of origin. It was originally domesticated in China and is still the most important legume crop throughout East Asia until today. It was first introduced to Europe and the US in the 18th century. During the 20th century it became a major crop in the Americas, and is currently the most widely cultivated legume crop of the world. In Central Europe (CEU), the area of soybean cultivation has been constantly increasing over the last decade and has revived the interest in breeding improved varieties for this production environment. This development is driven by the motivation to reduce dependencies from soy imports, to re-establish legumes in crop rotations for a more sustainable agriculture, and to mitigate the consequences of climate change by cultivating thermotolerant crops (BMEL 2016; European Soya Declaration 2017).

The adaptation of soybean to Central European production environments is an ongoing effort. So far, breeding research mainly focused on characterizing adaptive phenotypic variation of elite breeding germplasm by investigating the genetic basis of time to flowering and maturity (Kurasch et al. 2017). A portfolio of CEU cultivars that are classified into maturity groups (MGs) exists. In analogy to the system used in Northern America, MGs express the suitability of a variety for cultivation within a certain latitudinal range to ensure a full utilization of the vegetation period and timely maturation. Most cultivars suitable for production in Central Europe belong to the earliest MGs *000-0* and are mainly derived from North American material (Hahn and Würschum 2014). This narrow genetic basis (Gizlice et al. 1994) can limit long-term breeding progress and the utilization of genetic resources may alleviate this situation.

In addition to photoperiod, temperature and daily irradiation influence soybean development. Cold tolerance ensures that the vegetation period in high-latitude cold regions is fully exploited (Kurasch et al. 2017; Balko et al. 2014), but the trait has received little attention in soybean research so far. Flowering is the most susceptible developmental stage for low temperature, which causes flower and pod abscissions (Gass et al. 1996). Poor growth in early development and insufficient grain filling during pod development also reduce grain yield (Littlejohns and Tanner 1976). Phenotyping for cold tolerance, especially at flowering stage is laborious and a reliable exposure to cold stress requires controlled conditions, which limited the diversity of germplasm that has been evaluated for this trait (Funatsuki et al. 2004; Cober et al. 2013; Balko et al. 2014). Previous studies indicated a polygenic basis of cold tolerance and a potential overlap with the *E* series loci (Funatsuki et al. 2005) that are responsible for adaptation to photoperiod and soybean diffusion across latitudes (Jiang et al. 2014). Central European varieties exhibit variation in cold tolerance (Balko et al. 2014) and are used for improving cold tolerance in breeding of varieties for Northern Japan (Yamaguchi et al. 2015). Additional useful variation for cold tolerance and adaptation to high-latitude regions may be present in Asian landraces that are conserved in *ex situ* germplasm collections.

In this study, we present the development of two core collections of the USDA Soybean Germplasm Collection (Nelson 2011) with a focus on environmental adaptation to high-latitude cold regions to support the breeding of varieties for cultivation in Central Europe. Following the FIGS approach, we used environmental data characterizing the geographic origin of landraces and other germplasm in combination with current and future environmental conditions of the Central European *Target Population of Environments*. First, a precore collection consisting of accessions potentially adapted to CEU was formed based on environmental data. A scan for genetic differentiation between precore accessions and non-selected germplasm identified selection signals in genomic regions known to be involved in environmental adaptation and thus confirmed that FIGS enriched the precore for adaptive genetic variation. From the environmentally defined precore, we constructed genetically diverse, but substantially smaller core collections based on marker information. These may be used by researchers and breeders for comprehensive phenotypic characterization to facilitate trait discovery for Central European production environments. We show that a combination of FIGS and core collection methodology to define core subsets driven by targeted objectives will likely contribute to a more rapid and efficient utilization of soybean genetic resources.

## MATERIALS & METHODS

### Plant Material and the Donor Population of Environments

The *G. max* accessions originated from the USDA Soybean Germplasm Collection and only the *Introduced G. max* sub-collection (*N* > 17, 000) was considered, because it includes landraces (Nelson 2011; Song et al. 2015). Out of 6,180 accessions with georeferences we retained 5,886 Asian accessions by excluding material collected west of 60°E to remove accessions outside the original soybean cultivation range. The collection sites of the remaining accessions constitute the *Donor Population of Environments* (DPE) of Asian landraces, which may include potentially useful germplasm for CEU soybean breeding.

### Target Population of Environments

The *Target Population of Environments* (TPE) is the set of environments to which germplasm needs to be adapted for a successful cultivation. Our TPE consists of the soybean production environments in Central Europe (CEU) and includes Germany, Austria, Switzerland, Poland, Czech Republic and Slovakia. To characterize the TPE, we georeferenced locations that previously hosted soybean variety trials (Recknagel 2015; Kurasch et al. 2017). These served as proxies for the geographic extent of soybean cultivation in CEU since no detailed records regarding soybean cultivation exist for this region.

### Environmental Data

Based on the georeferences of the curated collection, environmental data was retrieved from the `WorldClim` (version 2) database (Fick and Hijmans 2017) at a resolution of 30 arc-seconds (≈ 1 km^2^). We used variables that are informative for the soybean cropping season in the northern hemisphere which lasts from May to September. Additionally, we modified the bioclimatic variables to render them informative for the soybean cropping season: Instead of the *annual mean temperature* (`BIO1`) we computed the *seasonal mean temperature* as the average mean temperature from May to September. Likewise we proceeded with `BIO2` to `BIO4`, `BIO6`, `BIO7` and `BIO12` to `BIO15`. `BIO8` to `BIO11` (*mean temperature of wettest / driest / warmest / coldest quarter*) were replaced by *mean temperatures of wettest / driest / warmest / coldest month* between May and September. `BIO16` to `BIO19` (*precipitation of wettest / driest / warmest / coldest quarter*) were replaced by monthly substitutes. An overview of the data used is provided in Table S1.

We also calculated monthly sums of *Crop Heat Units* (CHU) to quantify the accumulation of temperature over the growing season. CHUs are commonly used in Canada to rate the regional suitability of warm-season crop varieties with differing thermal requirements (Bootsma et al. 2007) and were also adopted by Central European soybean growers. Average daily CHU were computed from monthly averages of daily maximum and minimum temperatures according to Brown and Bootsma (1993):

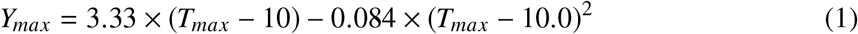

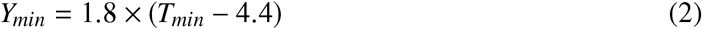

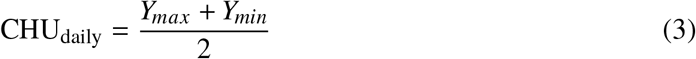

where *T_max_* and *T_min_* refer to the monthly averages of daily maximum and minimum temperatures, respectively. Negative *Y_max_* and *Y_min_* values were set to zero. A similar dataset was assembled for the TPE considering climate data for both current and future conditions. Climate projections for the year 2070 were retrieved from `WorldClim` version 1.4 (Hijmans et al. 2005) for the greenhouse gas scenario RCP8.5 as modeled by the *Community Climate System Model* (CCSM4.0). Available projections included average monthly climate data for minimum and maximum temperature, precipitation and the bioclimatic variables, and were used to compute derived variables as outlined above. Environmental qualities without projections for the year 2070 were substituted by their current equivalents to preserve the dimensional consistency of the datasets. Latitude information for the collection sites was included as additional environmental variable to account for the rather narrow adaptation of soybean germplasm to photoperiod.

### Environmental Characterization and Selection of Precore Accessions

Principal component analysis (PCA) was performed to summarize the high dimensional datasets of the DPE and the TPE separately using the `dudi.pca()` function from `R ade4` (Dray and Dufour 2007). Variables were mean centered and normalized. Correlations (i.e. the loadings) and squared loadings between the original variables and the principal components were used to quantify the amount of shared information and the contribution of single variables to the components. Environmental data for the TPE (current and projected for 2070 climate) was used to project Central European soybean growing environments as illustrative observations into the DPE’s multi-environmental space. The position of a collection site in the multivariate space was then used as ‘environmental profile’ associated with accessions of corresponding descent. Accessions with an environmental origin similar to environments of the TPE along the first two principal components were subsequently included in the precore to form a group of promising accessions with potential abiotic adaptation to the TPE.

### Genotypic Data

The USDA Soybean Germplasm Collection has been previously genotyped with the SoySNP50K array (Song et al. 2013; Song et al. 2015) and genotypes were downloaded from *SoyBase* (https://soybase.org/snps/; May 16, 2017) in the version mapped to the second *G. max* genome assembly “Glyma.Wm82.a2” (Schmutz et al. 2010). Non-chromosomal SNPs were removed from the genotype dataset and missing data was imputed using `BEAGLE` version 4.1 with default settings (Browning and Browning 2016).

### Phenotypic Data

Phenotypic data for the *Introduced G. max* sub-collection of the USDA Soybean Germplasm collection (Oliveira et al. 2010; Nelson 2011) was retrieved from https://npgsweb.ars-grin.gov/gringlobal/search.aspx. Data included qualitative and quantitative trait data like maturity group, 100 seed weight, seed yield, protein and oil contents. We used the quantitative trait data to assess changes in phenotypic performance and variance in the selection of the core collection. Homogeneity of variances in groups was tested using the modified robust Brown-Forsythe Levene-type test based on the absolute deviations from the median as implemented in `R lawstat` version 3.2 (Hui et al. 2008). Significance of differences of the means between groups was assessed with Welch’s unequal variances t-test implemented in `R` version 3.4.3 with Bonferroni correction for multiple testing.

### Core Sampling

Core sampling is based on reducing the number of available accessions by means of redundancy reduction through penalizing core subset solutions with too many too similar accessions. The large-scale genotyping of germplasm collections revealed the magnitude of identical and highly similar accessions in the USDA Soybean Germplasm Collection (Song et al. 2015) and other crops.

Core collections were assembled by analysing genotype data with `R Core Hunter` version 3.2 (http://www.corehunter.org/). All accessions that qualified after the environmental characterization and/or had a maturity group rating from 000 to I were considered for core sampling. We used `Core Hunter` in default execution mode, i.e. using the advanced parallel tempering search algorithm, and fixed the maximum number of steps without improvement to 100 (Beukelaer et al. 2017a; Beukelaer et al. 2017b) to efficiently select core subsets from the population of all possible cores (Thachuk et al. 2009). We explored different optimization strategies to identify the approach most suitable to our data and objectives: Sampling objectives regarding (1) optimization (i.e. maximization) of allelic diversity included the parameters expected heterozygosity and Shannon’s diversity index; (2) maximization of genetic distance based on the average entry-to-nearest-entry distance (Beukelaer et al. 2012) using the Modified Roger’s distance (MR) between accessions; (3) optimization of the auxiliary parameter allele coverage was performed the proportion of alleles retainable in core collections. For the definition of these measures the reader is referred to Thachuk et al. (2009). Core sizes of 5%, 10%, 15% and 20% of the input number of accessions were cross-evaluated for all three optimization objectives. Additionally, the simultaneous optimization of expected heterozygosity and MR was examined with varying weights to estimate the trade-off between both approaches. The measures expected heterozygosity, MR and allele coverage were also assessed for all sets used in this study to evaluate the success of the sampling process.

We also explored the effect of different sampling strategies on the subsequent utilization of the core collection: (1) Cores were sampled without any stratification; (2) they were sampled by means of classic stratification according to maturity, i.e. cores were sampled separately within three groups of accessions corresponding to maturity group ratings 000 to 0, I and II to X. A final core was then obtained by merging the result from these three groups; (3) a more refined 2-fold pseudostratification sampling strategy was devised and consisted in first sampling within accessions of maturity groups 000 to 0 and adopting the result of this first run as fixed in a second and final sampling run from all accessions, and (4) a 3-fold pseudostratification strategy first sampled within maturity groups 000 to 0, too. The second run then was limited to maturity groups 000 to I while fixing the result of the first run. In a third and final run, sampling was performed from all accessions while fixing the result from the second run. The general rationale behind all stratification strategies was primarily to retain accessions in the cores relative to their input group sizes by mitigating the effect of varying levels of genetic similarities within groups on the core sampling process. Table S4 provides a more detailed overview of the four core sampling strategies. For all reported cores the random seed was fixed to one prior to sampling for the sake of reproducibility of the core subset selection. To validate the stability of the results, re-sampling was performed for five additional independent runs with random seeds to rule out convergence effects.

### Screening for Signals of Adaptation

To identify genomic regions that reflect adaptive genetic differentiation between precore and non-selected accessions we used the basic model of `BAYPASS` (Gautier 2015) for allele count data. Only marker data for loci with minor allele frequencies ≥0.01 in the *Introduced G. max* subcollection was considered. `BAYPASS` calculates the *Xt X* statistic that may be seen as a SNP-specific *F_ST_* and accounts for the variance–covariance structure of the groups for population structure correction (Günther and Coop 2013). Significance of the observed *Xt X* signals was assessed from allele count data for 1,000 SNPs sampled randomly from a pseudo-observed dataset (POD). The POD was constructed with the R function `simulate.baypass()` based on the covariance matrix of population allele frequencies (Ω) and other properties of the original data (Gautier 2015). `BAYPASS` was then run on the simulated data to produce an empirical distribution of *Xt X* estimates and the 99% quantile value was used as threshold to distinguish selection from neutrality.

## RESULTS

### Curation of Collection Sites

Of 6,180 USDA soybean accessions with georeferenced collection sites and `WorldClim` data, 5,886 fell within our definition of the DPE. These originated from 699 unique collection sites, constituting an average contribution of ≈8 accessions per site. The distribution of accession numbers per collection site (Fig. S1) revealed several locations that contributed a large number (up to 619) of accessions to the collection, suggesting that multiple accessions from a region were aggregated into a single georeference. To purge the data set of accessions with unreliable geographic information, we checked locations with ≥20 accessions and found that georeferences were partly assigned based on provenance from cities, districts or even provinces (https://npgsweb.ars-grin.gov/gringlobal/search.aspx?). We removed 1,704 accessions (29%) from ten collection sites that were considered too large for a reliable inference of environmental adaptation (Fig. S1).

### Identification of Candidate Accessions for the TPE

By analyzing climate data from the geographic origin of accessions, we aimed at selecting accessions pre-adapted to cool and high latitude environments of Central Europe, and to identify the selective environmental factors acting during the soybean cropping season from May to September that characterize the DPE and TPE. Most environmental variables were strongly correlated with each other (Fig. S2 - S4). A PCA summarized 71.8% of the total environmental variation among collection sites in the first two principal components (PCs) (Fig. 1). Latitude and temperature based variables were strongly correlated with the first PC, while precipitation based variables related to the first and the second PC (Fig. S5 and S6). The first PC separated cooler from warmer regions of origin. To identify locations with accessions suitable for cultivation in Europe, we projected the European soybean cultivation environments onto the multivariate DPE to relate the TPE to the DPE (Polygons in Fig. 1). The projection shows that current CEU environments cluster in a narrow environmental range of less than 3,000 CHUs that includes a small proportion of provenances. Climate projections for 2070 indicate an average increase of more than 500 CHUs during the soybean cropping season in the TPE (Fig. S7). Even these conditions in the TPE exclude the majority of accessions because they originate from warmer regions with >3,500 CHUs. But the overlap of DPE and TPE with respect to temperature and other environmental variables increased and included North-Eastern China, Northern Japan, the Himalayas and Russia. The latter regions also contributed accessions from areas that appear to be highly unfavorable for soybean cultivation. To select potentially suitable accessions for cultivation in CEU, we included 123 environments with a position of ≥ 5 on the first PC (grey dashed line in Fig. 1) that provided 523 accessions with potential beneficial variation for abiotic adaptation to CEU (Fig. 2).

**Fig. 1.**
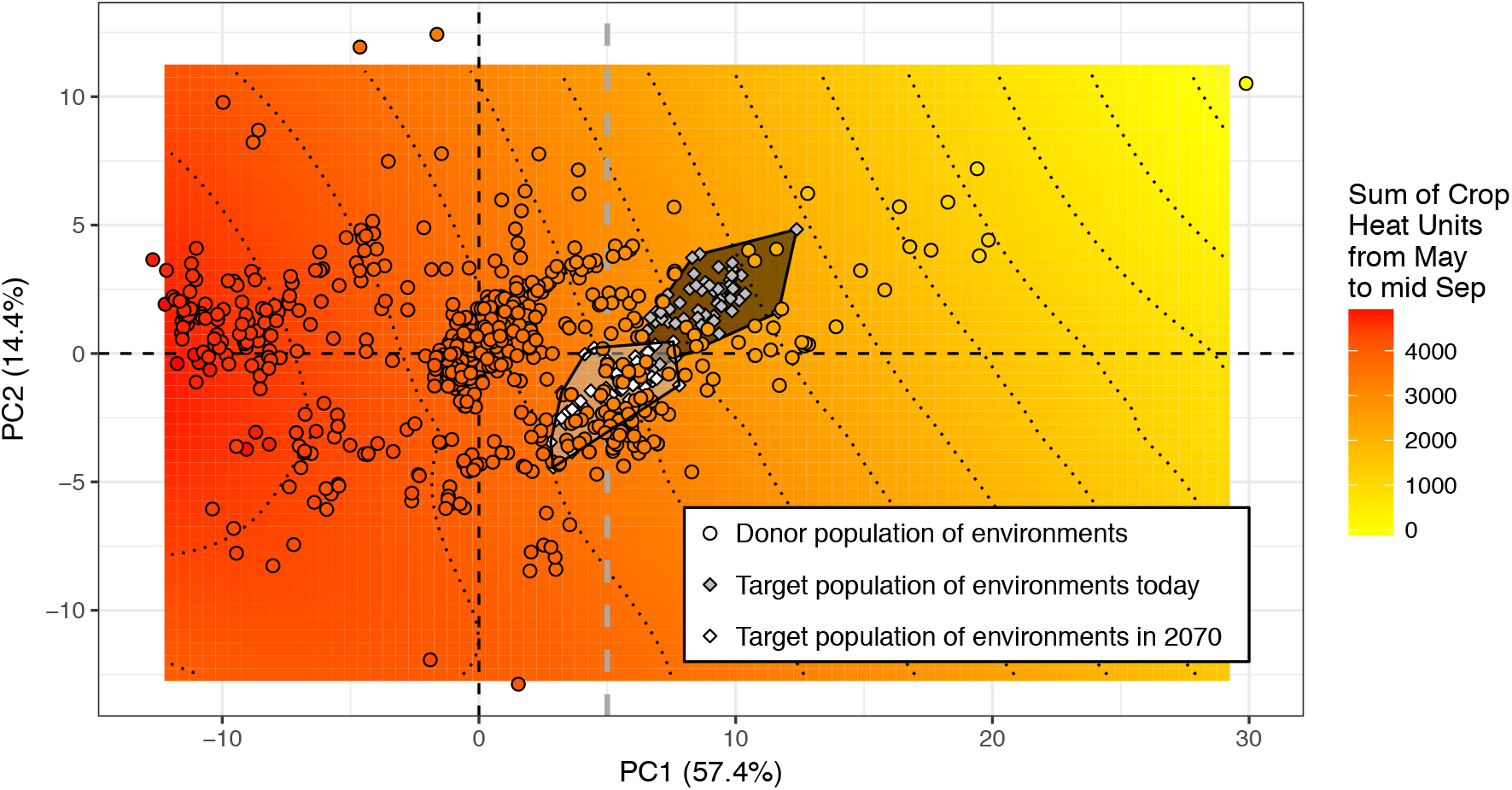
Principal component analysis of environmental variables obtained from *G. max* collection sites to characterize the donor population of environments (DPE). The color of collection sites corresponds to the sum of Crop Heat Units (CHUs) of the soybean cultivation season from May to mid September as a site-specific estimate of available temperatures. Contour lines represent bins of 400 CHUs. Current (grey) and future (white) climatic conditions in Central European soybean growing environments are included as illustrative observations. The environmental scopes of Central European scenarios (today and in 2070) are indicated with polygons.

**Fig. 2.**
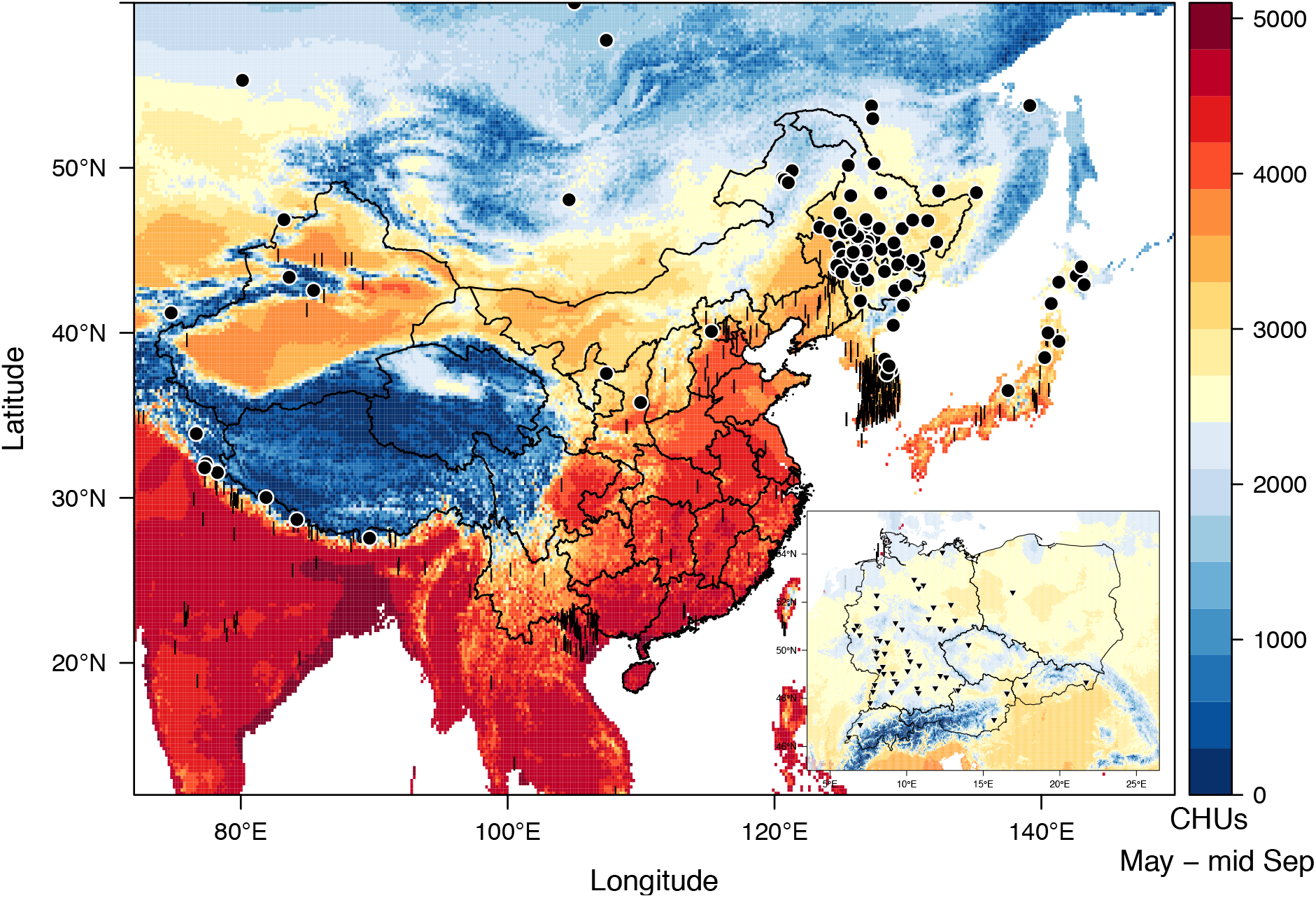
Collection sites of *G. max* germplasm in the *Donor Population of Environments*: Filled dots represent sites that were identified as donors of candidate accessions for CEU. Vertical lines represent sites which were not selected to contribute germplasm. Colouring represents estimates of available termperatures during the soybean season in Crop Heat Units. The inset shows the *Target Population of Environments* (current situation) with triangles indicating the extent of soybean cultivation in CEU based on soybean variety testing locations.

Only ≈20% of accessions from the USDA *Introduced G. max* collection were georeferenced and included in the above analysis of environmental variation in the DPE. However, the complete collection was categorized for maturity group, which is a proxy for the photo-thermal requirements of accessions and georeferenced accessions show a strong correlation between CHUs and their maturity group classification (Fig. S11). Accessions from early maturity groups originate from environments that resemble CEU e.g. in terms of temperature while late maturing accessions originate from warmer regions. We also included accessions without georeferences and no environmental data, but were classified as early maturing (maturity groups 000-I) and potentially suitable for cultivation in CEU. By combining accessions selected based on environmental data or maturity group classifications, we constructed an ‘environmental precore collection’ of 3,663 accessions (Tables 3 and 4).

### Genomic footprints of environmental adaptation

To test whether soybean accessions are locally adapted to their original cultivation environment, we searched for genomic regions with allele frequency differences between the 3,663 environmental precore accessions and the remaining 13,354 USDA *Introduced G. max* accessions. The `BAYPASS` program identified a total of 52 genomic regions with a stronger genetic differentiation (i.e. higher *XtX* values) than expected given the population structure (Fig. 3). These regions likely reflect genetic differentiation due to local adaption because they harbor several well characterized maturity genes of soybean, including *E1 - E3* on chromosomes 6, 10 and 19, which are three out of four cloned genes known to be major determinants of soybean adaptation to latitudinal zones (Jiang et al. 2014). On chromosome 19, a strongly differentiated region harbors *Dt1*, which is an ortholog of the *Arabidopsis thaliana TFL1* gene. In soybean, this gene controls the agronomically important trait indeterminacy and affects the length of flowering and reproductive period, which impacts plant height and maturity (Liu et al. 2010). The strongest differentiation occurs in a genomic region that contain the *Pdh1* and *GmFT2a* genes. The *pdh1* allele conveys shattering resistance and has been hypothesized crucial for soybean adaptation to Northern and semi-arid environments as opposed to humid South Asian environments where selection on pod dehiscence was more relaxed (Funatsuki et al. 2014). *GmFT2a* is a homolog of *Arabidopsis FLOWERING LOCUS T* (Sun et al. 2011) that has been identified as soybean maturity locus *E9* (Zhao et al. 2016). Additional significantly differentiated genomic regions harbor genes for which a role in abiotic adaptation of soybean has been demonstrated or postulated (Tab. 1 and S2). Phenotypic traits controlled by these genes include adaptation to photoperiod, heat, drought and cold stress. The limited marker density of the SoySNP50K array did not allow a fine-scaled mapping of differentiation signals, but in summary provides evidence that the environmental precore collection reflects such an adaptation. It also illustrates the possibility for mining the precore accessions for further traits that might not yet have been introduced into CEU breeding but could benefit soybean cultivation in cold and high-latitude environments. We used loci known to control environmental adaptation (i.e. *Dt1*, *E1-E3*, *E9* and *Pdh1*) to test whether genetic differentiation caused by local adaptation is better identified by environmental parameters or maturity group ratings. In general, genome scans comparing early maturing accessions (MG 000 to I) to late material (MG II to X) and comparing accessions of CEU-like origin to accessions of non CEU-like origin produced very similar results (Fig. S16). One exception was the *E1* locus region that exhibited a weaker differentiation signal in the `BAYPASS` run with accessions grouped solely based on environmental data as accessions of CEU-like origin were not necessarily of early maturity. This may indicate imprecise or incorrect georeferencing, which puts late maturing accessions in the wrong environments. On the other hand, these accessions might have acquired unique adaptation strategies that would not be identified by solely focusing on early maturing accessions. Therefore we decided to retain the respective accessions in the precore.

**Fig. 3.**
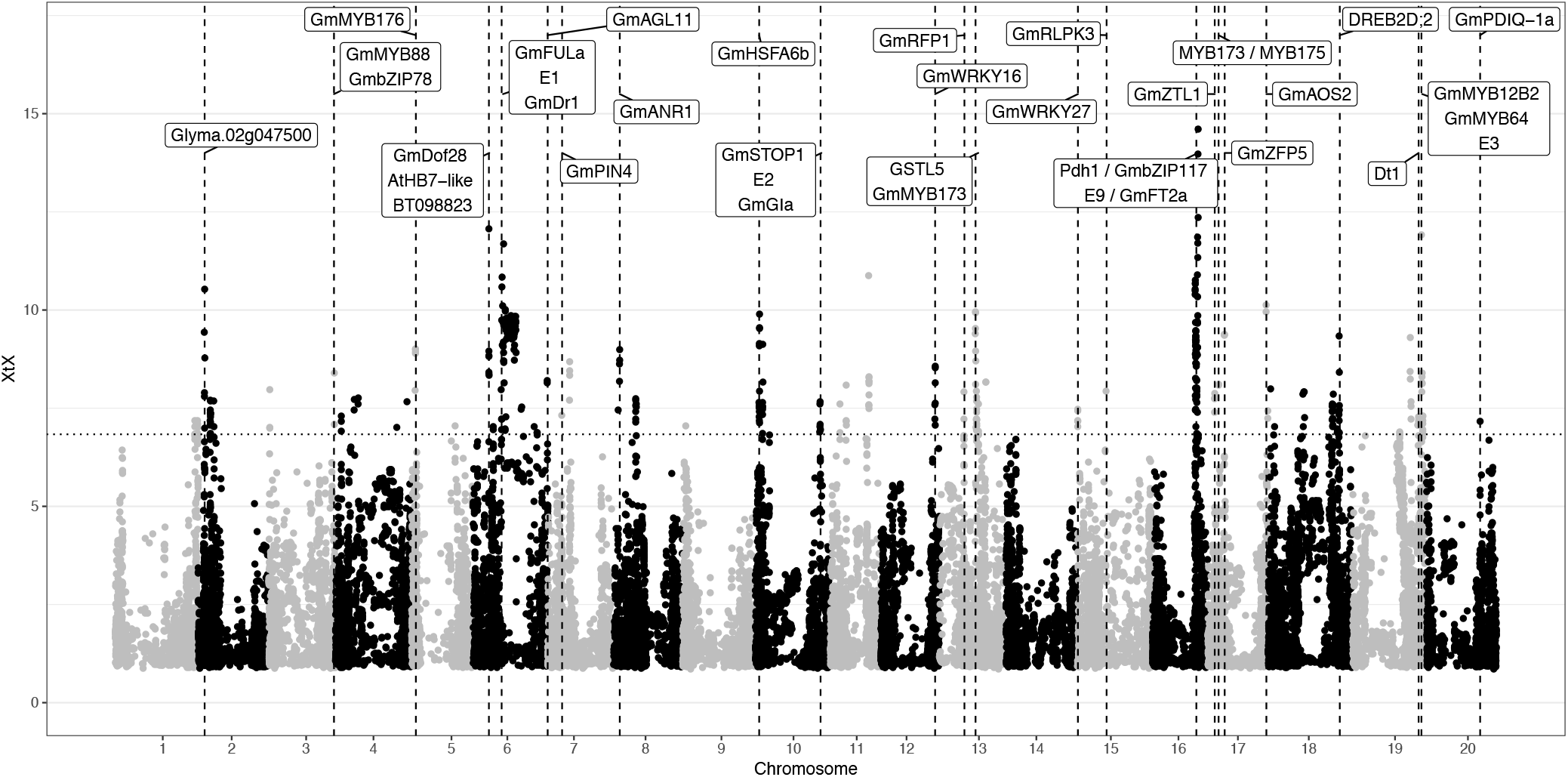
Genome-wide differentiation between precore and non-candidate accessions estimated with `BAYPASS`. Significantly differentiated regions are located above the dotted horizontal line which represents the POD 99% quantile of *XtX* values. Labels indicate genes that are located within significantly differentiated regions and known or hypothesized to be involved in abiotic adaptation and which are located within significantly differentiated regions. Chromosome-level plots are provided in Fig. S12 - S15. Candidate genes and their function are listed in Tab. 1 and Tab. S2.

**TABLE 1.**
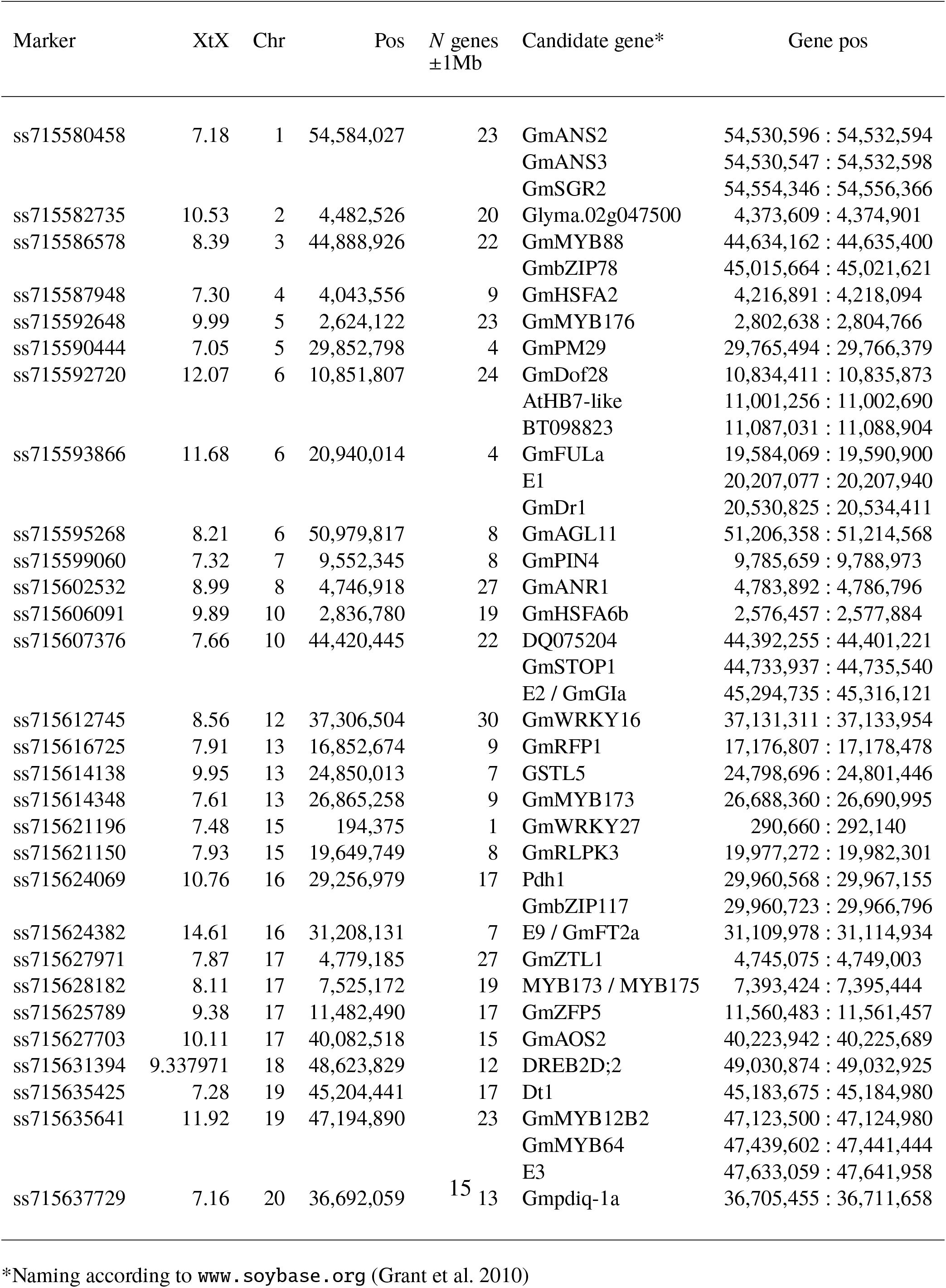
Overview of genomic regions significantly differentiated between the environmental precore and remaining accessions of the USDA *Introduced G. max* collection with proximity to known or hypothetical genes for abiotic adaptation. Gene functions are provided in Tab. S2.

### Construction of core collections

Since the environmentally defined precore comprised 3,663 accessions, we constructed core collections comprising 5% and 10% of the precore accessions to obtain manageable subsets for further evaluation by breeders and researchers.

#### Evaluation of Core Subset Optimization Strategies

We compared strategies that maximized allelic diversity within core collection accessions or maximized genetic distance between core entries, as well as a joint maximization of both parameters. As shown before (Thachuk et al. 2009), a combination of both parameters revealed a trade-off because the maximization of one parameter reduced the other (Fig. S17). Therefore, maximizing each parameter separately is the best approach for a given sampling objective (Fig. 4).

**Fig. 4.**
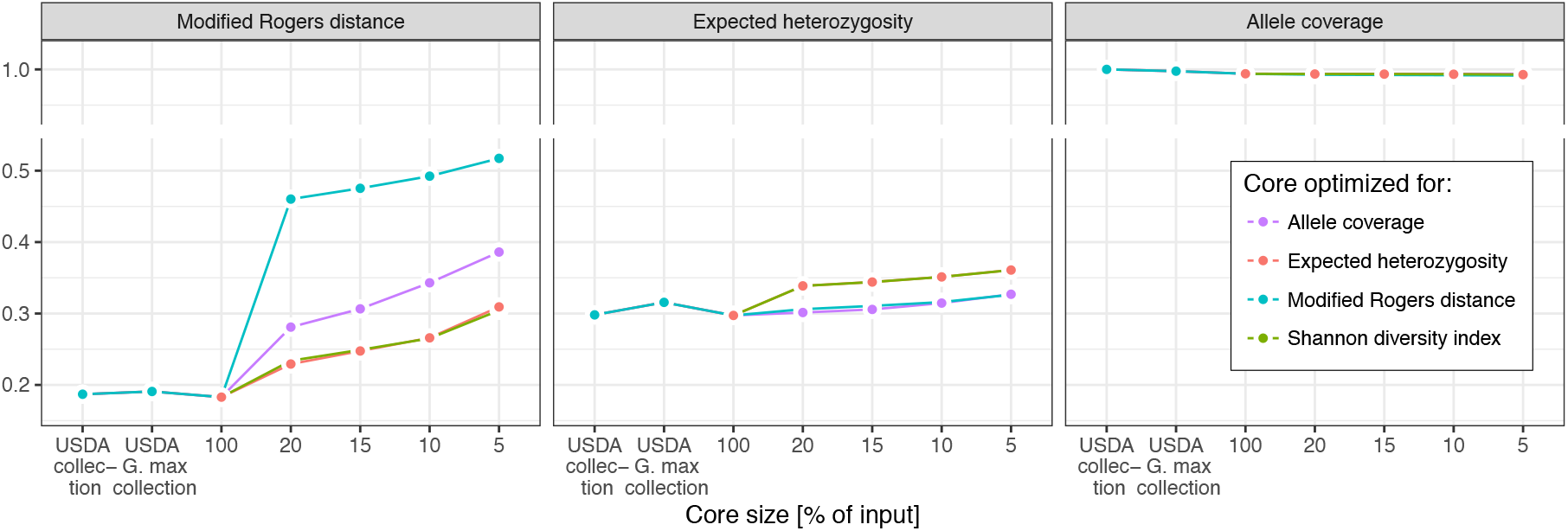
Genetic distance and allelic diversity preserved in cores of varying size with varying optimization objectives. Core size ‘100’ denotes the 3,663 precore accessions. Values on the y-axis refer to the respective facet label and all variables are defined in [0,1]. Core subset optimization based on Modified Rogers distance was most effective in forming cores consisting of distinct accessions.

Importantly, both approaches retained >99% of SNP alleles in all core collections. The loss of <1% of alleles resulted from the removal of *G. soja* specific alleles and the environmental stratification (Fig. S18). We compared two measures of allelic diversity, *H_exp_* and Shannon’s diversity index. Both gave very similar results with respect to diversity within and distance between accessions, and were increased in core collections (Fig. 4). The genetic distance between accessions increased more strongly in core collections than allelic diversity and this behavior was independent of the optimization strategy. For example, in the most successful scenario the average minimum Modified Rogers distance increased 1.5-fold in the 5% core compared to the precore whereas *H_exp_* differed only marginally (Fig. 4). Our estimates of allelic diversity (Table S3) might show an ascertainment bias caused by the design the SoySNP50K array if it includes SNPs that were preselected for high minor allele frequencies to guarantee high informativeness (Song et al. 2013). This most likely contributed to the modest increase of *H_exp_* in core subset optimization (Malomane et al. 2018). By contrast, core subset optimization based on Modified Rogers distance purged highly similar accessions from subsets and was effective in forming cores consisting of distinct accessions (Fig. 4). We therefore based the core subset optimization exclusively on the maximization of genetic distance between core accessions.

#### Evaluation of Core Subset Sampling Strategies

The construction of core collections resulted in an underrepresentation of early maturing accessions (Fig. 5A and S19) because average redundancy levels were higher in early than within late maturing accessions (Fig. S20). Since early maturity is an important trait for CEU soybean breeding we explored sampling strategies based on a stratification by maturity group designation that favored the selection of early accessions. Any stratification prior to optimization represents a trade-off between accepting higher levels of redundancy in exchange for gaining improved representation in the stratification criterion. The best relative representation across all maturity groups was therefore observed for subsets sampled according to a classic stratification strategy (Fig. 5A) of independent sampling within three groups of accessions and aggregating selected accessions into a final core (Table S4). But it also resulted in the lowest level of realized genetic distance between accessions (Fig. 5B) and reflects that accessions in one subcore can be similar to accessions in another, which reduces the overall diversity of the complete set. To increase genetic distance we adopted two pseudostratification sampling strategies that allowed to coordinate sub-core subset selection in a consecutive, step-wise fashion that ensured the selection of complementary sub-cores (Table S4). Sampling with pseudostratification resulted in cores that in terms of genetic dissimilarity between accessions were comparable to the no stratification benchmark. The 2-fold pseudostratification however failed in ensuring a good level of representation for accessions of maturity group *I*, presumably because accessions of this group on average are more similar to accessions of maturity groups *000 - 0* than to later accessions. The 3-fold pseudostratification resulted in a high average minimum genetic distance between accessions in the final set and also maintained acceptable levels of representation over all maturity groups. We therefore used the 3-fold pseudostratification to construct the final core collections with 5% and 10% of individuals from the precore collection.

**Fig. 5.**
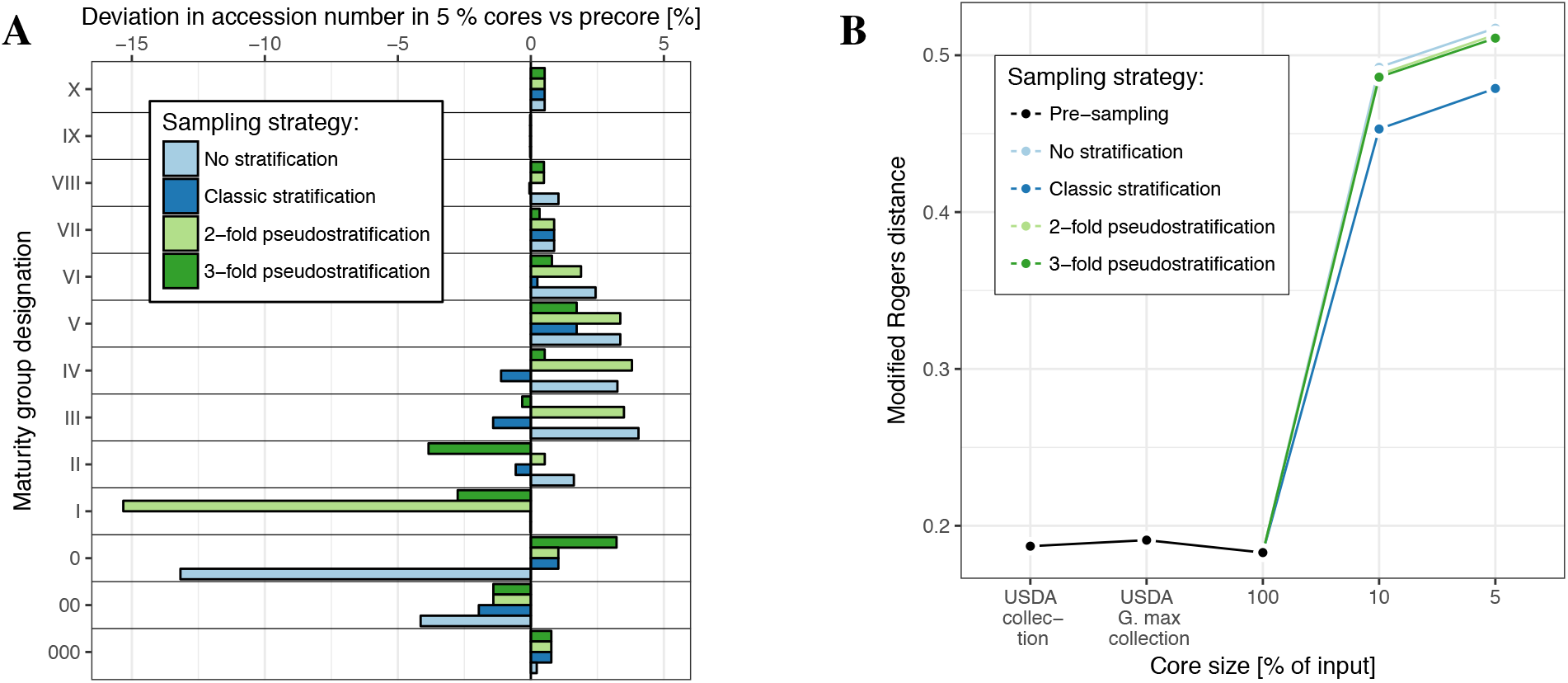
**A**: Evaluation of different core sampling strategies with regards to the preservation of MG fractions in 5% cores relative to the precore. **B**: Increase in genetic distance between core accessions measured by Modified Roger’s distance. Among different sampling strategies, the 3-fold pseudostratification^*^ showed good results in terms of favouring early maturing material while maintaining high average minimum genetic distance between accessions. ^*^Consecutive sampling within groups of maturity categories *000 - 0*, *000 - I*, *000 - X* with fixation of already picked accessions (Tab. S4).

#### Further Efects of Core Sampling: Shifts in Germplasm Composition and Allele Frequencies

The reduction of the number of accessions through our environmental stratification and the selection of core subsets maximized for genetic diversity was naturally accompanied by changes in the geographic and genomic composition of the respective accession population at each stage of the core formation process (Fig. 2, 6 and S21). The reduction of genetic redundancy among core entries furthermore resulted in genome wide changes in population-wise allele frequencies (Fig. S22). These mainly affected loci with low minor allele frequencies in the environmental precore by elevating the frequencies of the respective alleles in the cores (Fig. 8). Changes in MAFs between the 3,663 precore accessions and the core collection entries however never exceeded 0.25 (Fig. 22) and the groups showed a comparable distribution along the first two components of a PCA (Fig. 7). Also the genetic basis of the presumed environmental adaptation was unaffected by the core formation process and well preserved in the cores as can be observed in a comparison of genetic differentiation between core, precore and non-precore accessions (Fig. S16).

**Fig. 6.**
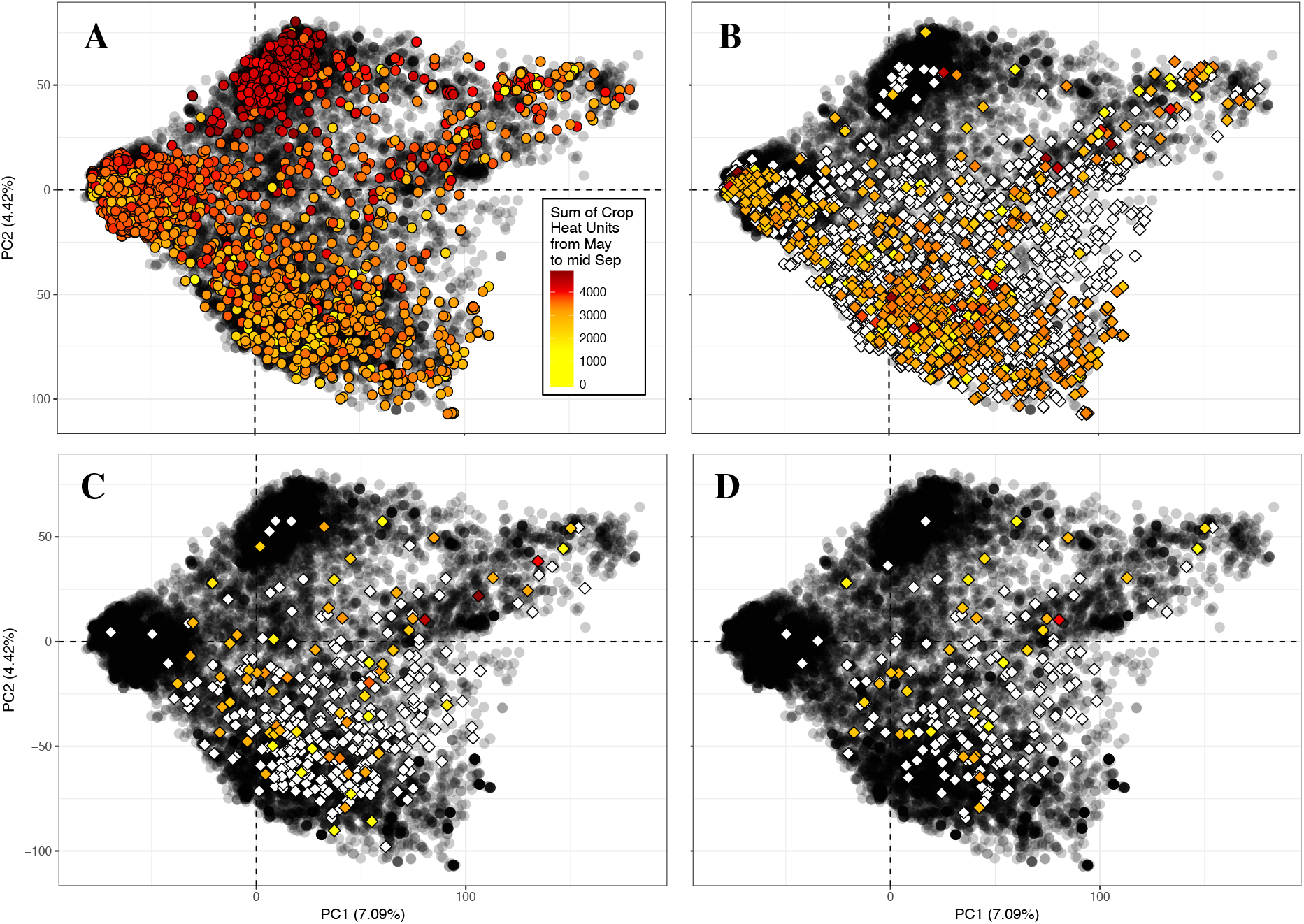
Principal component analysis summarizing the genetic structure in > 17.000 *G. max* accessions conserved in the *Introduced G. max* subcollection of the USDA Soybean Germplasm Collection. Each dot represents one accession. **A**: Accessions with collection site information available in their passport data are highlighted according to temperature supply at origin. **B**: 3,663 accessions selected for the environmental precore based on environmental data (coloured) and / or early maturity group ratings (white) are highlighted. **C**: 366 entries of the 10% core are highlighted. **D**: 183 entries of the 5% core are highlighted.

**Fig. 7.**
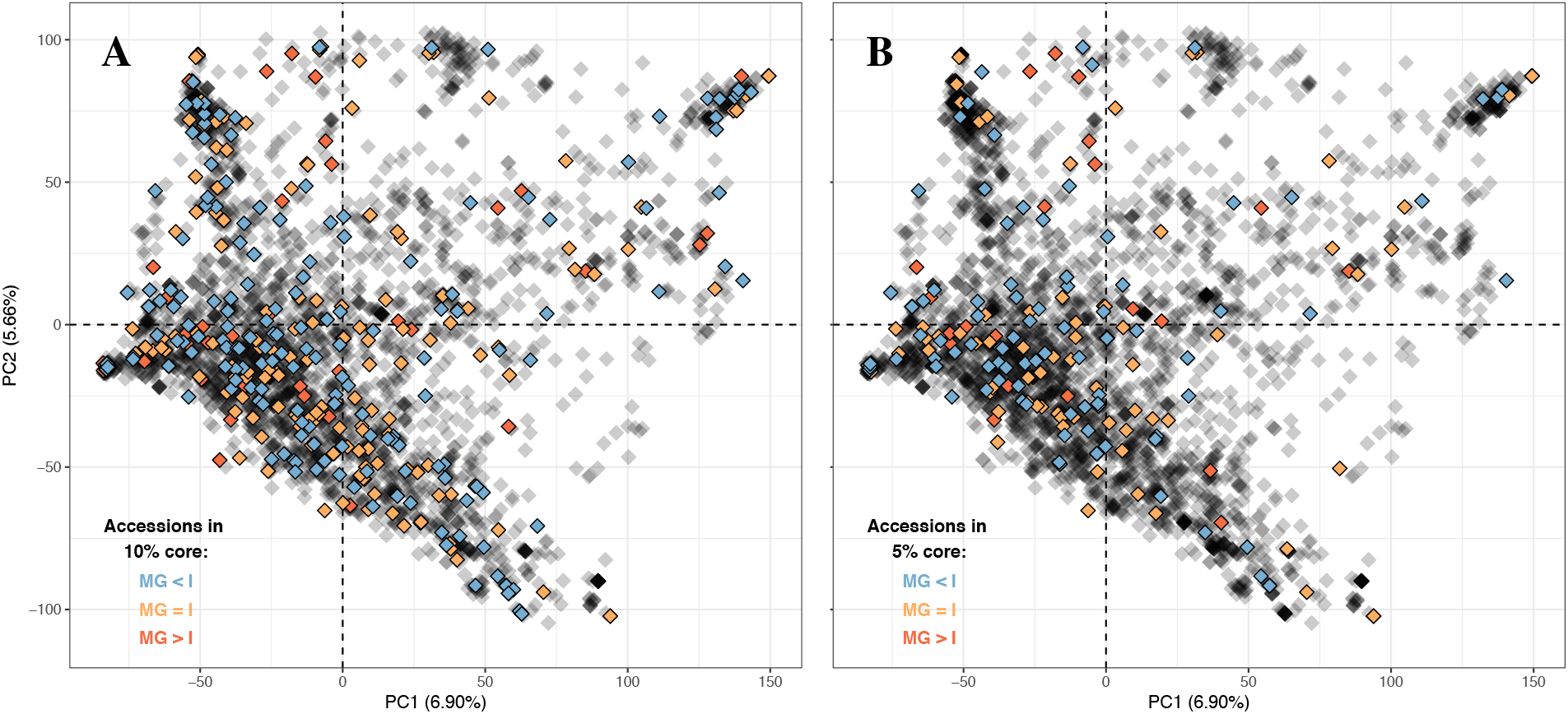
Principal component analysis summarizing the genetic structure among 3,663 precore accessions shows an even spatial distribution of the core entries: core accessions are highlighted according to maturity group ratings. **A**: 366 entries of the 10% core. **B**: 183 entries of the 5% core.

**Fig. 8.**
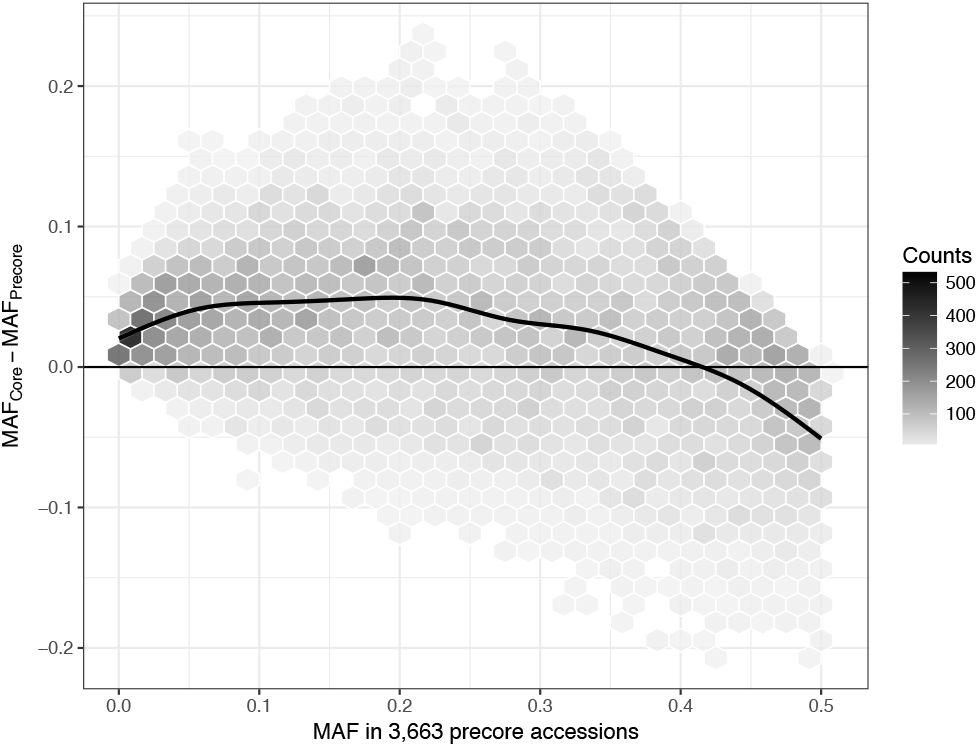
Pairwise comparison of MAFs in the precore versus the MAF change from precore to 5% core: Core subset optimization resulted in genome wide MAF changes through enrichment of low frequency variants in the core. Fig. S22 provides a more detailed overview on MAF changes.

#### Robustness of Core Sampling

With 3,663 precore accessions, 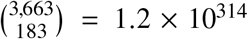 core subsets with 5% (*n* = 183) and 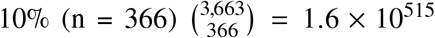 accessions are possible. To find the best core, stochastic algorithms to approximate the optimal solution as implemented in `Core Hunter` need to be used. Independent runs of these search algorithms may produce different results, but using a fixed seed to define the starting point ensures the reproducibility of the optimization solution as long as the termination criteria are kept constant. All results presented so far regarded core subset sampling solutions derived from optimization runs with a fixed seed of one. We verified the stability of our results from runs with fixed seeds by performing re-sampling with random seeds for five additional runs. Average levels of genetic distances between accessions in these runs were highly similar to the previous runs. The intersection of all 5% cores included the same 82 out of 183 (45%) and 223 out of 366 10%-core accessions (61%). Only a small proportion of accessions (9% / 5% in 5%/10% cores, respectively) was restricted to a single core subset and seemingly interchangeable through genetically similar entries (Table 2). Core subsets differing in sampling intensities were comparable on an individual level too, with 134 accessions of the 5% core having been also selected for the 10% core solution.

**TABLE 2.**
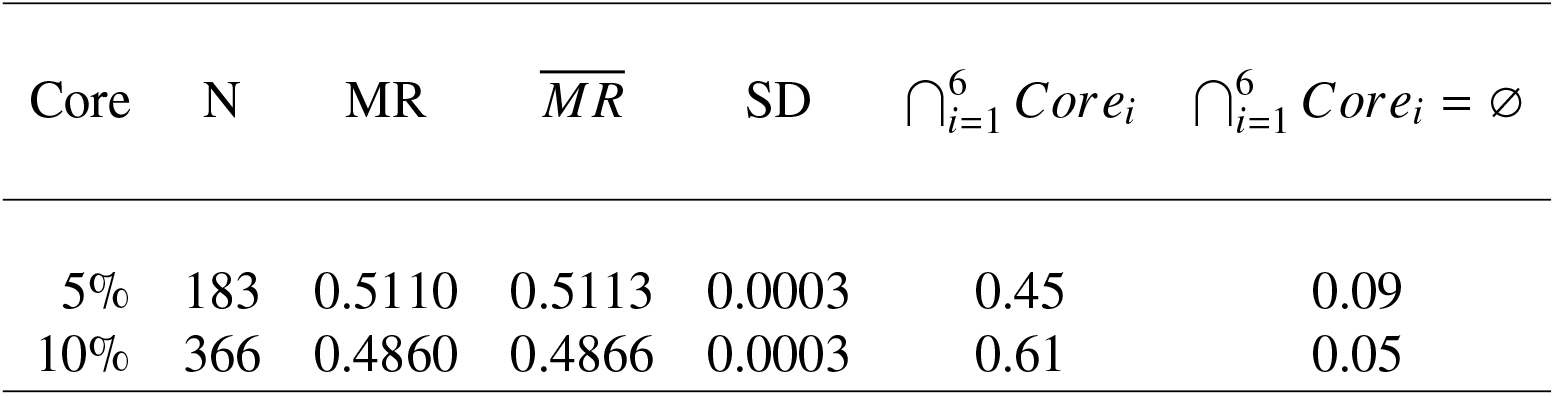
Comparison of core collection sampling alternatives: Sampling intensity, core size, averaged minimum Modified Rogers distance between core accessions sampled with fixed seed, mean and standard deviation of averaged minimum Modified Rogers distance between core accessions of all sampled cores (including cores sampled with random seed), proportion of accessions that jointly occur in all sampled cores, proportion of accessions private to one core.

| Core | N | MR | $\overline{MR}$ | SD | $\bigcap_{i=1}^6 Core_i$ | $\bigcap_{i=1}^6 Core_i = \emptyset$ |
| --- | --- | --- | --- | --- | --- | --- |
| 5% | 183 | 0.5110 | 0.5113 | 0.0003 | 0.45 | 0.09 |
| 10% | 366 | 0.4860 | 0.4866 | 0.0003 | 0.61 | 0.05 |

We followed Odong et al. (2013) and investigated how the formation of core collections depends on the marker set used by randomly dividing our marker set in a training set and an evaluation set of equal size (split-half analysis). The training set was used to sample cores with varying sampling intensities and the average minimum genetic distance between core entries was subsequently evaluated independently based on both marker sets and based on all available markers. Estimates of genetic distance were highly similar in all comparisons of the same sampling intensity and did not differ between marker sets (Δ*MR* 0.00032, data not shown). This suggests that our SNP marker set was sufficiently dense to warrant for the robust estimation of genetic distance and the presented core solutions can thus be considered stable within the disparities discussed above.

#### Evaluation of phenotypic properties

We assessed changes in major phenotypic characteristics of accession sets throughout the core collection formation process, i.e. from the *Introduced G. max* subcollection over the environmental precore to the actual core subsets. The largest phenotypic changes were caused by the environmental stratification that removed most accessions of maturity group ≥ *II* and therefore the majority of the original collection (Table 3). As a consequence, variation in phenotypic traits of the precore and core subsets differed substantially from the *Introduced G. max* collection while the smaller core collections preserved the phenotypic variation of the precore (Table 4). Regarding the average phenotypic performance, seed weight was reduced in the cores compared to the precore while the seed yields did not differ significantly and seed components (protein and oil content) varied partially. A reduction in seed weight may result from the marker-based selection of more diverse exotic accessions instead of more recently adapted material with larger seeds. Cores sampled with random seeds produced similar results (Table S5).

**TABLE 3.**
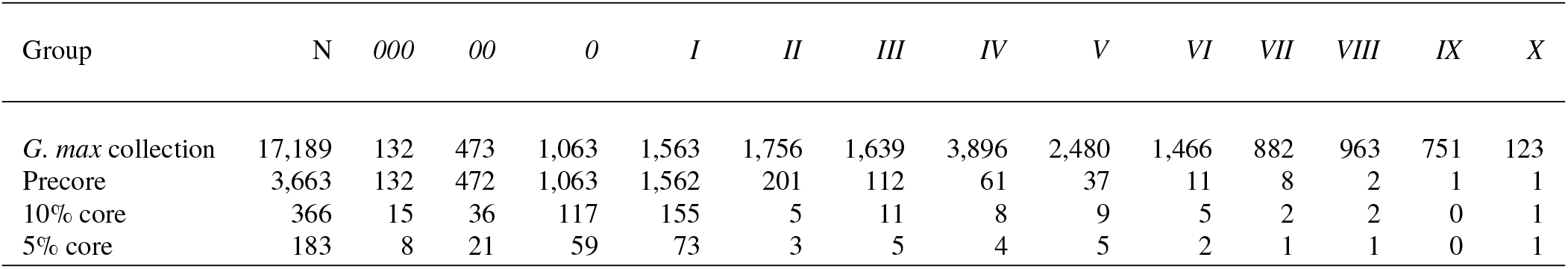
Composition of the *Introduced G. max* sub-collection with regards to maturity group ratings and the shift towards earlier maturity in precore accessions and core entries.

| Group | N | 000 | 00 | 0 | I | II | III | IV | V | VI | VII | VIII | IX | X |
| --- | --- | --- | --- | --- | --- | --- | --- | --- | --- | --- | --- | --- | --- | --- |
| *G. max* collection | 17,189 | 132 | 473 | 1,063 | 1,563 | 1,756 | 1,639 | 3,896 | 2,480 | 1,466 | 882 | 963 | 751 | 123 |
| Precore | 3,663 | 132 | 472 | 1,063 | 1,562 | 201 | 112 | 61 | 37 | 11 | 8 | 2 | 1 | 1 |
| 10% core | 366 | 15 | 36 | 117 | 155 | 5 | 11 | 8 | 9 | 5 | 2 | 2 | 0 | 1 |
| 5% core | 183 | 8 | 21 | 59 | 73 | 3 | 5 | 4 | 5 | 2 | 1 | 1 | 0 | 1 |

**TABLE 4.** Selected phenotypic properties of the *Introduced G. max* sub-collection, the environmental precore and the core collections. Differences between groups are reported as significant at a p-value < 0.05.

| Group | N | Yield<br>[Mg/ha] | Var | Seed weight<br>[cg/seed] | Var | Protein<br>[%] | Var | Oil<br>[%] | Var |
| --- | --- | --- | --- | --- | --- | --- | --- | --- | --- |
| *G. max* collection | 17,189 | 1.954<sup>a</sup> | 0.503<sup>a</sup> | 15.31<sup>a</sup> | 29.53<sup>a</sup> | 44.33<sup>a</sup> | 7.98<sup>a</sup> | 17.81<sup>a</sup> | 4.36<sup>a</sup> |
| Precore | 3,663 | 2.283<sup>b</sup> | 0.479<sup>b</sup> | 15.63<sup>b</sup> | 16.41<sup>b</sup> | 42.69<sup>b</sup> | 7.23<sup>b</sup> | 18.88<sup>b</sup> | 3.04<sup>b</sup> |
| 10% core | 366 | 2.262<sup>b</sup> | 0.547<sup>ab</sup> | 14.67<sup>c</sup> | 13.36<sup>b</sup> | 43.18<sup>c</sup> | 6.88<sup>b</sup> | 18.53<sup>c</sup> | 3.30<sup>b</sup> |
| 5% core | 183 | 2.149<sup>b</sup> | 0.578<sup>ab</sup> | 13.96<sup>c</sup> | 16.25<sup>b</sup> | 43.30<sup>bc</sup> | 10.05<sup>ab</sup> | 18.21<sup>ac</sup> | 4.77<sup>ab</sup> |

## DISCUSSION

### Creation of Core Collections

By combining the *Focused Identification of Germplasm Strategy* (FIGS) with core collection methodology, we identified subsets of genebank accessions from a large soybean germplasm collection (N > 17,000) that are likely pre-adapted to cultivation in Central Europe while simultaneously conserving a high level of genetic diversity. To establish these core collections, we first defined a precore of 3,663 accessions with potential abiotic adaptation to CEU based on environmental data. This precore was subsequently reduced to two core collections of 183 and 366 accessions (5% and 10% of the precore, please consult the supplementary information for further core accession details) by limiting genetic redundancies among core entries. We quantified the success of this approach at each step of the process and found strong evidence for the enrichment of adaptive genetic variation for abiotic adaptation to high-latitude cold regions throughout the genome (Fig. 3, S12 to S16, Tab. 1 and Tab. S2).

### Limitations and Evidence for Enrichment of Environmental Adaptation

By inferring local adaptation from environmental data we followed FIGS methodology that is based on the assumption that landraces adapted to the environment in which they were originally cultivated by natural and artificial selection (Street et al. 2008; Sanders et al. 2013). Knowledge about the geographic origin of germplasm is therefore a key parameter in the successful application of FIGS for abiotic traits. For our implementation of FIGS in the USDA Soybean Germplasm Collection, the origin of accessions was approximated by the georeference for the respective collection sites recorded in the passport data. This information was only available for parts of the collection and in many cases seemed to be an approximate reconstruction of the historic collection sites, sometimes with limited accuracy. Inaccurate georeference information lead to false associations of accessions with environmental data and even though we removed the least reliable collection site - origin misspecifications, this certainly led to the inclusion of some non-adapted accessions into the environmental precore. Another limitation concerns the granularity of the environmental data: The spatial resolution and categorical scope of WorldClim data is sufficiently detailed, but the temporal resolution is restricted to monthly averages. Since records on local cropping practices such as the period of cultivation of landraces in their native environment are lacking too, a detailed reconstruction of the agro-ecological conditions that influenced local adaptation was not possible. Missing georeferences for the majority of accessions were even more limiting, which required us to use maturity group assignments of these accessions as sole proxy for environmental adaptation. Although field trials for the maturity group classification were conducted in North America, we consider using these information suitable for this purpose because of the high heritability of this trait (Xavier et al. 2018).

To test whether the selection of the precore accessions led to an enrichment of abiotic adaptation to CEU environments, we conducted a selection scan for genetic differentiation between precore accessions and non-precore accessions. Although the marker density of the SoySNP50K array was too low to pinpoint differentiation at the level of single genes, we found evidence that the selection based on climate data and maturity group assignments targets genomic regions involved in abiotic adaptation and enriches genetic variants which are advantageous for cultivation in CEU in the pre-core. Highly differentiated regions mainly indicated genomic regions that harbor well-known genes responsible for photoperiodic adaptation in soybean. This result is expected because the selection of adapted accessions was partially based on maturity groups that are conditioned by photoperiod genes. In addition, also new and promising candidate genes for adaptation to abiotic factors like heat, drought and cold stress were highlighted (Tab. 1 and Tab. S2). Most importantly, these signals of adaptation were preserved in the core collections (Fig. S16) which indicates that the genetic variation responsible for this presumed adaptation can be introgressed from core accessions into CEU elite breeding germplasm.

### Towards a Characterization of Core Collections

To unlock the full potential of the core accessions, long term dedication of breeders and researchers for the detailed genotypic and phenotypic characterization of the cores will be necessary. A list of core entries is included as supplementary information for further reference. Environmental adaptation is complex with possibly manifold ways to adapt to the same stress which in a real world scenario will not be present as a single factor but as a combination of multiple factors. Therefore, multi-location field trials will be necessary to accurately estimate the level of abiotic adaptation to CEU environments, complemented by trials with controlled conditions in which the most promising accessions can be confronted with selected stresses. It should be noted that no accession is likely to outcompete current CEU varieties per se, but may contribute to improve breeding germplasm. Detailed phenotyping will identify beneficial variation in landraces for use in soybean prebreeding. The increasing availability of high-throughput aerial and field-based phenomics technologies holds great potential in this respect (Furbank and Tester 2011; Araus and Cairns 2014).

The efficient introduction of beneficial abiotic adaptation into breeding germplasm via marker assisted selection or genome editing requires the identification of causal genetic variants. The allele frequency of causal variants is a major determinant for the detection power in association-mapping panels (Spencer et al. 2009) and variants with MAF below 5% are usually not considered because of their low statistical power. Our optimization strategy for the selection of core accessions resulted in elevated MAF for rare alleles (Fig. 8), which will aid in the discovery of rare traits using modern phenotyping approaches and provides an increased power to map causal variants despite the fairly small core collection sizes. Accessions that harbor favorable alleles can be subsequently included in large multiparent populations for genetic fine-mapping and gene identification (Cockram and Mackay 2018).

### Core Collections for a Future Climate

The time span required for the development of new varieties requires long-term planning in an era of climate change. We therefore selected precore accessions not only using current climate parameters of the TPE, but also considered climate projections for the year 2070, in which lower precipitation and higher temperatures during the soybean cropping season are expected for CEU (Fig. S7 and S8). These projections suggest that future varieties may not require cold adaptation, but will have to be highly drought tolerant. It is reasonable to argue that both types of adaptation will remain relevant because climate estimates are based on long-term averages, whereas short-term extreme events during the cropping season are frequently limiting crop yields. The future climate is expected to become more volatile and extreme events more frequent, which will include low temperatures in the early and high temperatures in the later growing season. The improvement of crop abiotic adaptation is therefore pivotal to equip agricultural systems with crop varieties for the future. The novel combination of FIGS and core collection methodology is a promising strategy to assemble relevant objective driven core collections and to increase the efficiency of germplasm characterization. Both will aid the identification of alleles for abiotic stress tolerance in order to complement modern breeding germplasm. Upon discovery, targeted crosses and combinations of marker-assisted selection and speed breeding or genome engineering can be employed to incorporate beneficial alleles into modern backgrounds in real-time (Varshney et al. 2018; Watson et al. 2018). Genomic selection has emerged as a strategy to predict complex traits also in genebank populations (Jarquin et al. 2016; Schnable et al. 2016), but its successful application is challenging in genetically diverse germplasm collections that are frequently lacking phenotypic characterization data for traits of interest (Langridge and Waugh 2019). Here, upon their sufficient characterization, objective driven core collections from FIGS selected germplasm groups have the potential to complement prediction efforts in genebank germplasm for abiotic adaptation by serving as training and validation populations.

## AUTHOR CONTRIBUTION STATEMENT

MH and KS designed the study, MH analyzed the data and wrote the first version of the manuscript. MH and KS wrote the final version of the manuscript.

## ACKNOWLEDGEMENTS

The authors thank the United States Department of Agriculture (USDA) and the USDA Soybean Germplasm Collection for their open data and free distribution of germplasm policies that made this work possible. This work was funded by BMEL Project Sojagenopath (FKZ 2814EPS012).

## CONFLICT OF INTEREST STATEMENT

On behalf of all authors, the corresponding author states that there is no conflict of interest.

