## Supplementary tables and figures for "Combining Focused Identification of Germplasm and Core Collection Strategies to Identify Genebank Accessions for Central European Soybean Breeding"

**Tab. S 1.** Overview of climate variables that were used to characterize the donor population of environments as well as the current and future target population of environments.

| Variable | Measure |
|---|---|
| lat | Latitude (°) |
| tavg | Average monthly mean temperature for months May - Sep (°C) |
| tmin | Average monthly minimum temperature for months May - Sep (°C) |
| tmax | Average monthly maximum temperature for months May - Sep (°C) |
| sCHU_May2MidSep | Sum of daily Crop Heat Units from May - mid Sep |
| sCHU_Month | Monthly sum of Crop Heat Units for months May - Sep |
| prec | Average monthly precipitation for months May - Sep (mm) |
| srad | Average monthly solar radiation for months May - Sep ($MJ\ m^{-2}$)* |
| vapr | Average monthly vapour pressure for months May - Sep (kPa)* |
| wind | Average monthly wind speed for months May - Sep ($m\ s^{-1}$)* |
| BIO1_May2Sep | Seasonal mean temperature (mean of tavg for months May - Sep) |
| BIO2_May2Sep | Mean diurnal range (mean of tmax - tmin for months May - Sep) |
| BIO3_May2Sep | Isothermality (BIO2_May2Sep / BIO7_May2Sep) * 100) |
| BIO4_May2Sep | Temperature seasonality (standard deviation of tavg * 100 for months May - Sep) |
| BIO5 | tmax of warmest month |
| BIO6_May2Sep | tmin of coldes month for months May - Sep |
| BIO7_May2Sep | Temperature seasonal range (BIO5 - BIO6_May2Sep) |
| BIO8_May2Sep | tavg of wettest month for months May - Sep |
| BIO9_May2Sep | tavg of driest month for months May - Sep |
| BIO10_May2Sep | tavg of warmest month for months May - Sep |
| BIO11_May2Sep | tavg of coldest month for months May - Sep |
| BIO12_May2Sep | Seasonal precipitation (sum of prec for months May - Sep) |
| BIO13_May2Sep | prec of wettest month for months May - Sep |
| BIO14_May2Sep | prec of driest month for months May - Sep |
| BIO15_May2Sep | Precipitation seasonality (Coefficient of variation of prec for months May - Sep) |
| BIO18_May2Sep | prec of warmest month for months May - Sep |
| BIO19_May2Sep | prec of coldest month for months May - Sep |

Note: BIO16 and BIO17 (prec of wettest quarter and prec of driest quarter) were not assessed for the 5-month soybean cropping season. BIO13_May2Sep and BIO14_May2Sep provide the equivalent at a monthly resolution.
*For these variables only estimates for the current climate were available.

**Tab. S 2.** Overview of gene function for genes listed in Tab. 1.

| Gene | (Potential) Function in abiotic adaptation (taken from Grant et al. (2010), if not cited otherwise) |
|---|---|
| GmANS2 | Anthocyanidin synthase 2: anthocyanin pigments are plant secondary metabolites that have a variety of ecophysiological functions including protection from abiotic stresses (Kovinich et al. 2012) |
| GmANS3 | Anthocyanidin synthase 3: anthocyanin pigments are plant secondary metabolites that have a variety of ecophysiological functions including protection from abiotic stresses (Kovinich et al. 2012) |
| GmSGR2 | Senescence-inducible chloroplast stay-green protein 2 (Park et al. 2007) |
| Glyma.02g047500 | Gene model for cold shock domain containing proteins (Sasaki and Imai 2012) |
| GmMYB88 | *G. max* MYB transcription factors function in plant growth, developmental metabolism and stress responses and have been shown to e.g. enhance drought and cold tolerance in *At* (Su et al. 2014) |
| GmbZIP78 | GmbZIP78 is a negative regulator of ABA signaling and functions in salt and freezing tolerance (Liao et al. 2008) |
| GmHSFA2 | *G. max* heat shock transcription factor 2 |
| GmMYB176 | *G. max* MYB transcription factors function in plant growth, developmental metabolism and stress responses and have been shown to e.g. enhance drought and cold tolerance in *At* (Su et al. 2014) |
| GmPM29 | *G. max* seed maturation protein |
| GmDof28 | Dof transcription factors are associated with many plant-specific physiological processes including responses to abiotic stress (Wang et al. 2017b) |
| AtHB7-like | Probable transcription activator that may act as growth regulator in response to water deficit (Olsson et al. 2004) |
| BT098823 | Heat shock transcription factor |
| GmFULa | *At* FRUITFULL homolog (Bemer et al. 2017) |
| E1 | *G. max* maturity gene (Bernard 1971) |
| GmDr1 | TATA box binding protein involved in the repression of transcription (Song et al. 2002) |
| GmAGL11 | Promotion of flowering and maturity (Zeng et al. 2018) |
| GmPIN4 | PIN genes have been shown to be involved in soybean response to abiotic stress (Wang et al. 2015) |
| GmANR1 | Anthocyanidin reductase 1: anthocyanin pigments are plant secondary metabolites that have a variety of ecophysiological functions including protection from abiotic stresses (Kovinich et al. 2012) |
| GmHSFA6b | *G. max* heat shock transcription factor 6b |
| DQ075204 | Chorismate synthase, involved in aromatic amino acid synthesis |
| GmSTOP1 | *At* STOP1 homolog, involved in tolerance to acidic soils in *At* (Iuchi et al. 2014) |
| E2 / GmGIa | *G. max* maturity gene (Bernard 1971; Watanabe et al. 2011) |
| GmWRKY16 | *G. max* WRKY transcription factors have been shown to e.g. confer differential tolerance to abiotic stresses in *At* (Zhou et al. 2008) |
| GmRFP1 | RING-type E3 ubiquitin ligase, up-regulation by ABA and salt stress, down-regulation by cold and drought treatments (Du et al. 2010) |
| GSTL5 | Glutathione S-Transferases have important functions in the response to environmental conditions (McGonigle et al. 2000) |
| GmMYB173 | *G. max* MYB transcription factors function in plant growth, developmental metabolism and stress responses and have been shown to e.g. enhance drought and cold tolerance in *At* (Su et al. 2014) |
| GmWRKY27 | *G. max* WRKY transcription factors have been shown to e.g. confer differential tolerance to abiotic stresses in *At* (Zhou et al. 2008) |
| GmRLPK3 | Receptor-like kinase 3: expression and phylogenetic analysis suggest involvement in regulating soybean leaf senescence and stress responses (Ma et al. 2006) |
| Pdh1 | Pod dehiscence 1: *pdh1* conveys shattering resistance, especially important in northern, semi-arid environments (Funatsuki et al. 2014) |
| GmbZIP117 | bZIP-transcription factors have been shown to regulate abiotic stress responses in soybean (Liao et al. 2008) |
| E9 / GmFT2a | *G. max* maturity gene (Sun et al. 2011; Zhao et al. 2016) |
| GmZTL1 | Soybean homolog of *At* ZEITLUPE1 (Somers et al. 2000) |
| MYB173 / MYB175 | *G. max* MYB transcription factors function in plant growth, developmental metabolism and stress responses and have been shown to e.g. enhance drought and cold tolerance in *At* |
| GmZFP5 | Zinc finger proteins are involved in response to different environmental stresses in plants and a soybean ZFP has been shown to enhance cold tolerance in *At* (Yu et al. 2014) |
| GmAOS2 | Allene oxide synthase 2: involved in jasmonic acid synthesis which functions in regulation of responses to abiotic and biotic stresses as well as plant growth and development (Kongrit et al. 2007) |
| DREB2D;2 | *G. max* dehydration-responsive element-binding protein 2 family members have been shown to improve abiotic stress tolerance in *At* (Mizoi et al. 2013) |
| Dt1 | Ortholog of *At* TERMINAL FLOWER1 (Liu et al. 2010) |
| GmMYB12B2 | *G. max* MYB transcription factors function in plant growth, developmental metabolism and stress responses and have been shown to e.g. enhance drought and cold tolerance in *At* (Su et al. 2014) |
| GmMYB64 | *G. max* MYB transcription factors function in plant growth, developmental metabolism and stress responses and have been shown to e.g. enhance drought and cold tolerance in *At* (Su et al. 2014) |
| E3 | *G. max* maturity gene (Buzzell 1971) |
| Gmpdiq-1a | Protein disulfide isomerases are involved in protein folding in the endoplasmatic reticulum (ER) (Iwasaki et al. 2009), disruption of proper folding can result in ER stress responses (Silva et al. 2015) |

**Tab. S 3.** Expected heterozygosity in the USDA Soybean Germplasm collection and several subcollections. Subcollections according to Nelson et al. (2011).

| Subcollection | $H_{exp}$ |
|---|---|
| Complete USDA collection | 0.3156362 |
| *G. soja* collection | 0.2861149 |
| *Introduced G. max* collection | 0.2980931 |
| Old US cultivars collection | 0.3084224 |

**Tab. S 4.** Overview of the explored core sampling strategies.

| Core sampling strategy | |
|---|---|
| No stratification | Core sampling is performed without any grouping |
| Classic stratification | Core sampling is performed in three groups:<br>1.1 MG's 000 - 0<br>1.2 MG I<br>1.3 MG's II - X<br>2. Final core is compiled (merged) from MG group cores 1.1 - 1.3 |
| 2-fold pseudo-stratification | 1. sampling within MGs 000 - 0<br>2. sampling within MGs 000 - X while fixing results from 1st |
| 3-fold pseudo-stratification | 1. sampling within MGs 000 - 0<br>2. sampling within MGs 000 - I while fixing results from 1.<br>3. sampling within MGs 000 - X while fixing results from 2. |

**Tab. S 5.** Selected phenotypic properties of CCs sampled with random seeds. No significant differences between groups were observed (p-value < 0.05).

| Group | N | $\overline{Yield}$ [Mg/ha] | Var | $\overline{Seedweight}$ [cg/seed] | Var | $\overline{Protein}$ [%] | Var | $\overline{Oil}$ [%] | Var |
|---|---|---|---|---|---|---|---|---|---|
| 10% core | 366 | 2.262 | 0.547 | 14.67 | 13.36 | 43.18 | 6.88 | 18.53 | 3.30 |
| R10.1 | | 2.265 | 0.546 | 14.70 | 14.08 | 43.09 | 7.42 | 18.58 | 3.30 |
| R10.2 | | 2.247 | 0.542 | 14.52 | 13.38 | 43.14 | 7.42 | 18.54 | 3.53 |
| R10.3 | | 2.240 | 0.549 | 14.53 | 12.95 | 43.08 | 7.85 | 18.58 | 3.36 |
| R10.4 | | 2.265 | 0.518 | 14.51 | 13.22 | 43.13 | 7.82 | 18.59 | 3.48 |
| R10.5 | | 2.259 | 0.536 | 14.67 | 13.99 | 43.11 | 7.47 | 18.56 | 3.38 |
| 5% core | 183 | 2.149 | 0.578 | 13.96 | 16.25 | 43.30 | 10.05 | 18.21 | 4.77 |
| R5.1 | | 2.156 | 0.625 | 13.90 | 17.35 | 43.48 | 9.46 | 18.09 | 5.19 |
| R5.2 | | 2.168 | 0.610 | 13.92 | 15.96 | 43.53 | 8.69 | 18.20 | 4.83 |
| R5.3 | | 2.138 | 0.579 | 13.93 | 15.49 | 43.23 | 10.98 | 18.27 | 4.58 |
| R5.4 | | 2.147 | 0.602 | 14.06 | 16.51 | 43.32 | 9.45 | 18.22 | 4.80 |
| R5.5 | | 2.103 | 0.587 | 13.72 | 16.66 | 43.34 | 9.74 | 18.20 | 5.10 |

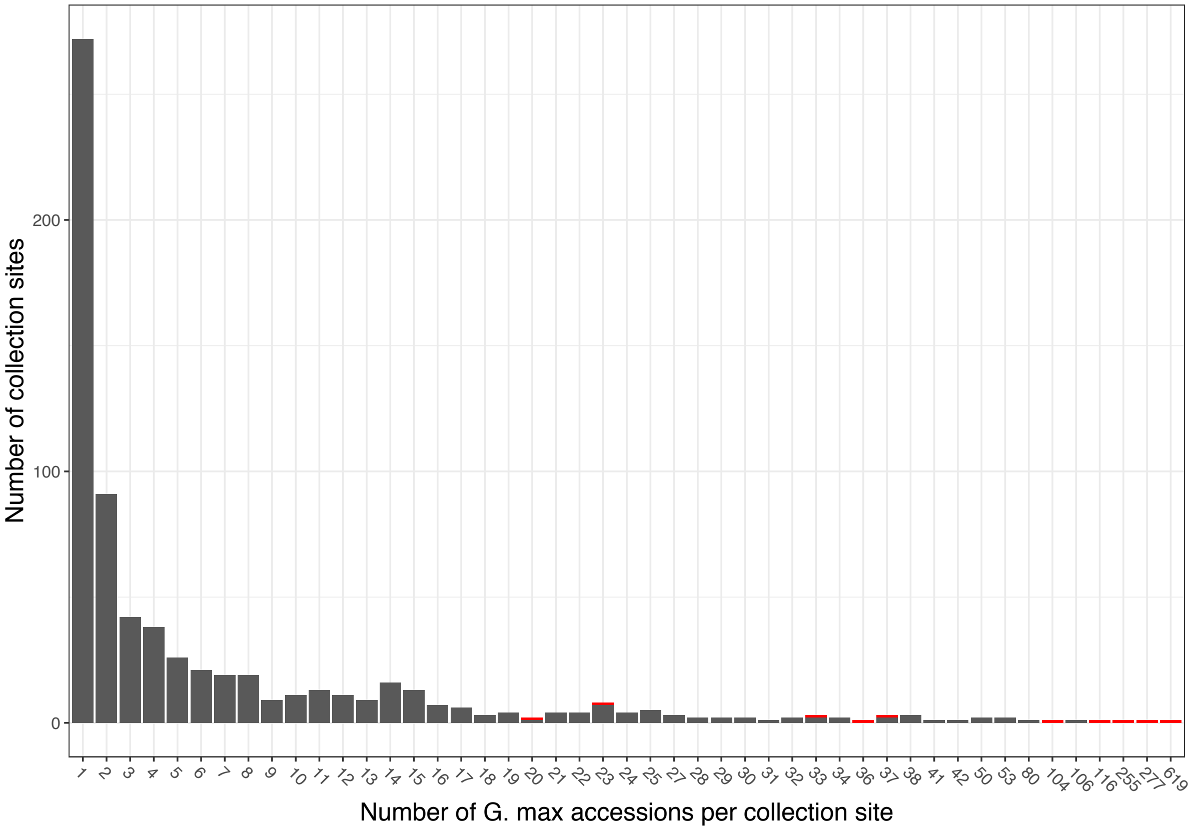


**Fig. S 1.** Collection site frequency spectrum summarizing the distribution of the number of *G. max* accessions that are recorded to have been collected at the same site (i.e. identical georeference) in the USDA Soybean Germplasm Collection. Red bars indicate collection sites that were identified as ambiguous, supposedly due to pooling of accessions from large areas into one georeference etc. Those were excluded from the subsequent environmental analysis. Examples of ambiguous sites of origin and excluded germplasm include 619 accessions form the Japanese regions Kanto and Tosan, 277 accessions from the Japanese region of Tohoku, 255 accessions from the Russian region of Primorsky and 116 accessions from the historic entity of Manchuria as well as further material of similarly unspecific origin. Early maturing accessions (MG 000 - I) from these sites were however reintroduced into the precore and thus considered for the final core solutions.

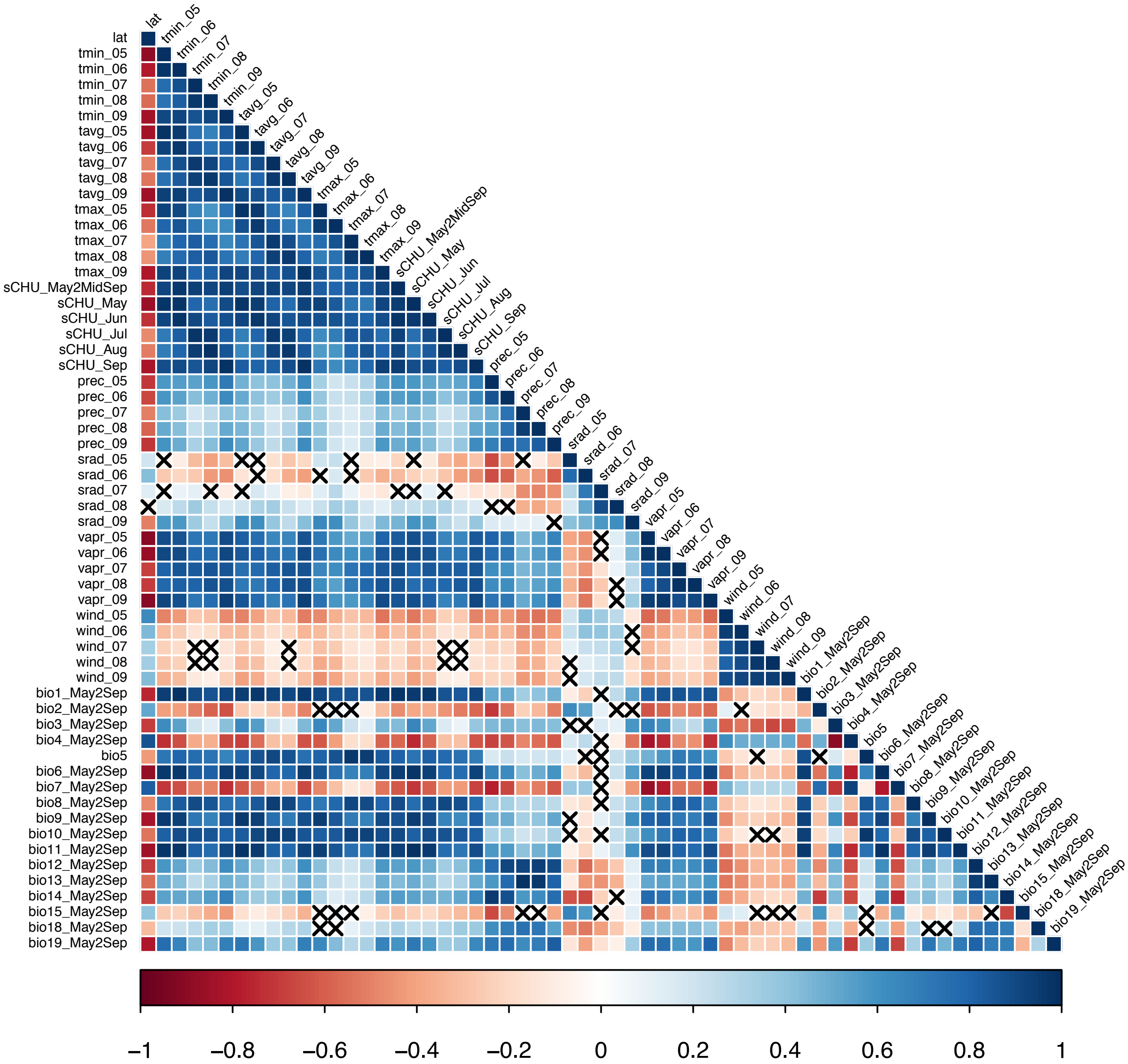


**Fig. S 2.** Correlations among environmental variables describing Asian collection sites of *G. max* accessions. Strength and direction of correlation is indicated by the colour key. Non-significant relationships ($p > 0.05$) are indicated by black crosses.

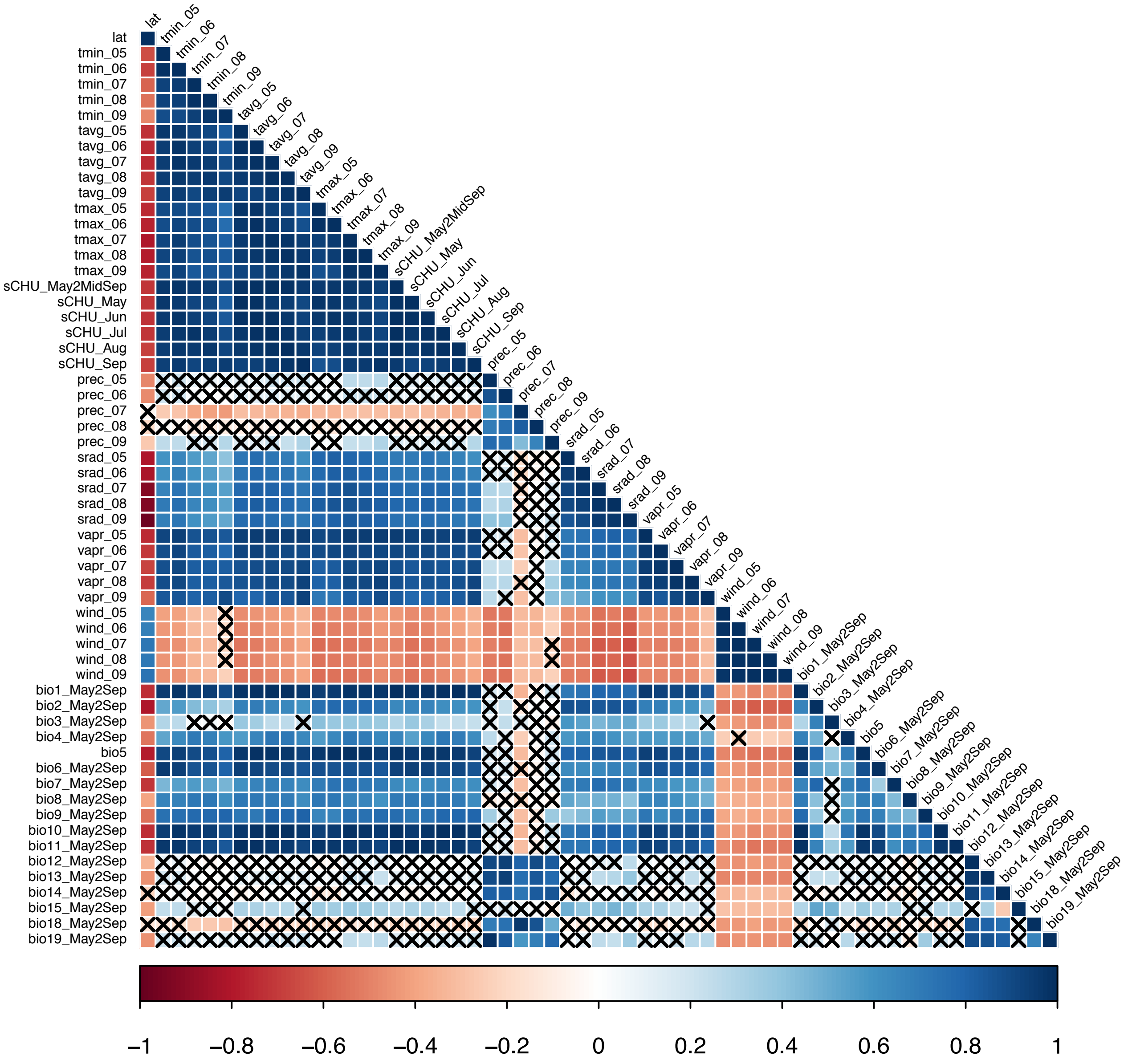


**Fig. S 3.** Correlations among environmental variables describing current European soybean growing environments. Strength and direction of correlation is indicated by the colour key. Non-significant relationships (p > 0.05) are indicated by black crosses.

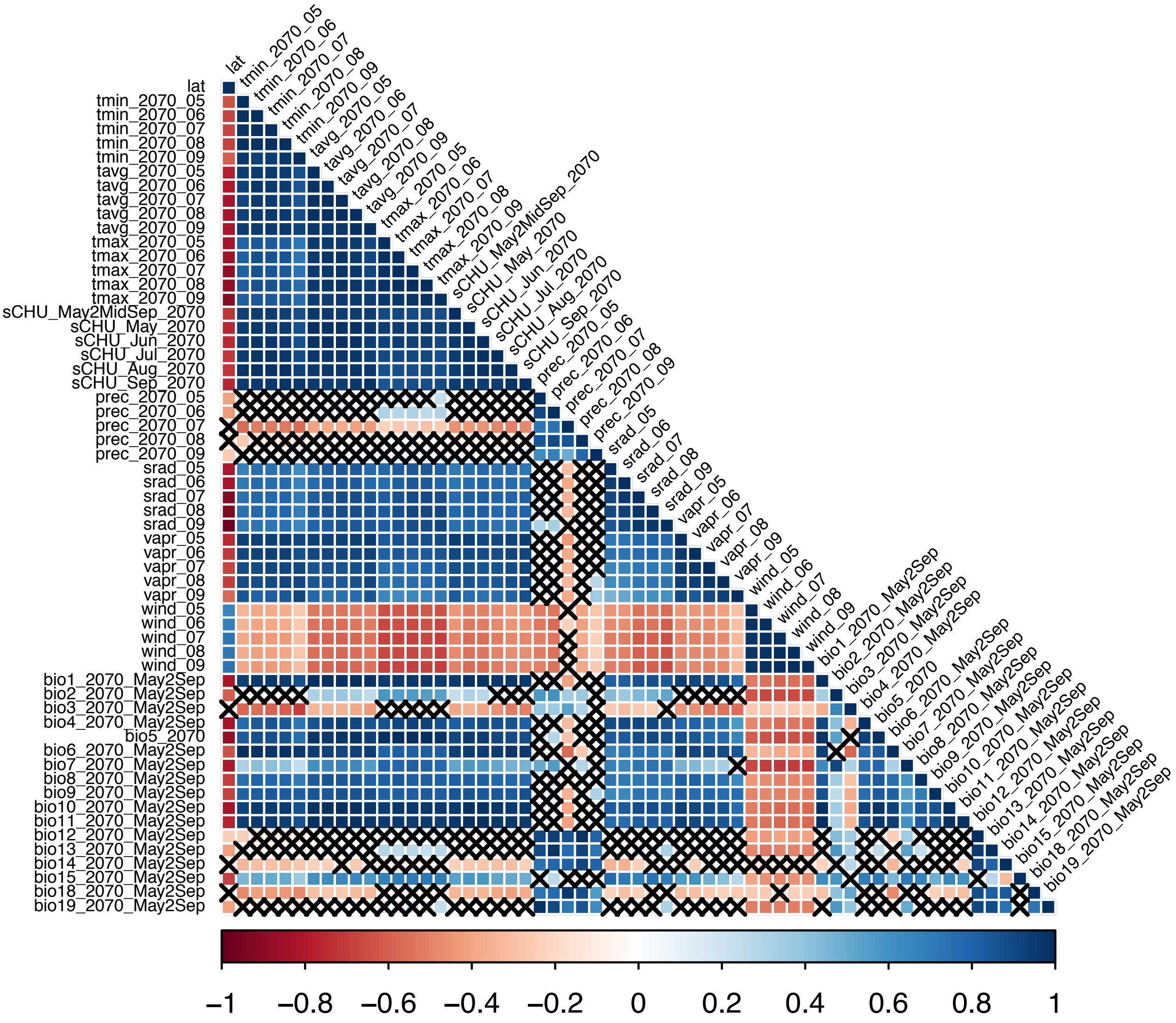


**Fig. S 4.** Correlations among environmental variables describing European soybean growing environments in 2070. Strength and direction of correlation is indicated by the colour key. Non-significant relationships (p > 0.05) are indicated by black crosses.

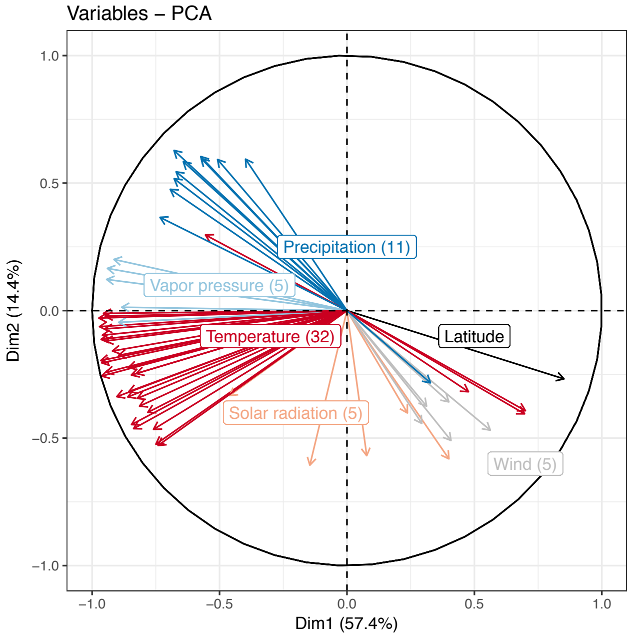


**Fig. S 5.** Correlation circle for the first two components of the PCA with environmental data characterizing Asian *G. max* collection sites. The correlation of a given variable with the first component can be read off from the x-axis and for the second component from the y-axis, respectively.

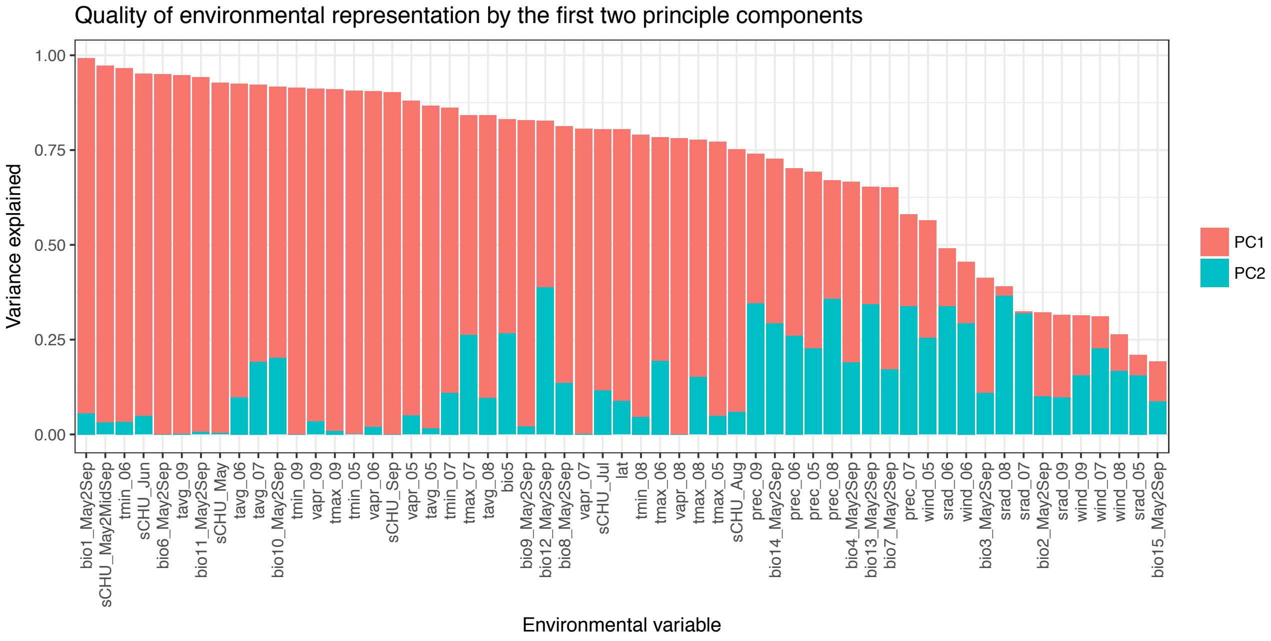


**Fig. S 6.** Quality of representation of original environmental variables characterizing Asian *G. max* collection sites jointly by the first two principal components.

PCA of current and future environmental conditions in European G. max growing areas

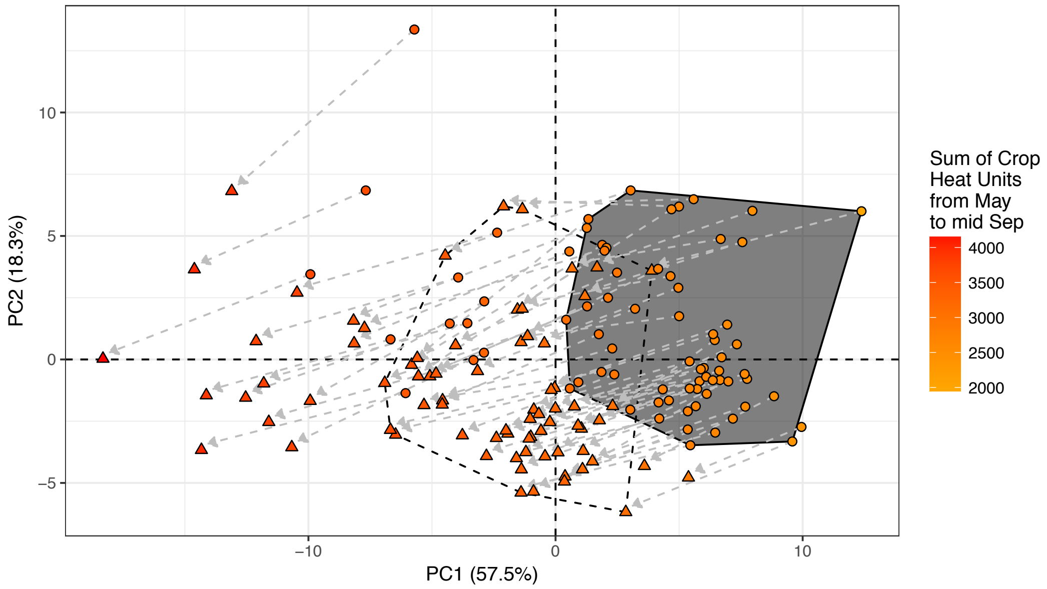


**Fig. S 7.** Distribution of Central (diamonds) and South (triangles) European soybean growing environments along the first two components of a PCA performed with environmental data characterizing the target population of environments. Colouring of sites was done according to the sum of Crop Heat Units for the approximate soybean season from May to mid September as a site specific estimate of available temperatures. Arrows indicate the projected change of conditions until 2070. Scopes of Central European scenarios (today and in 2070) are indicated with polygons.

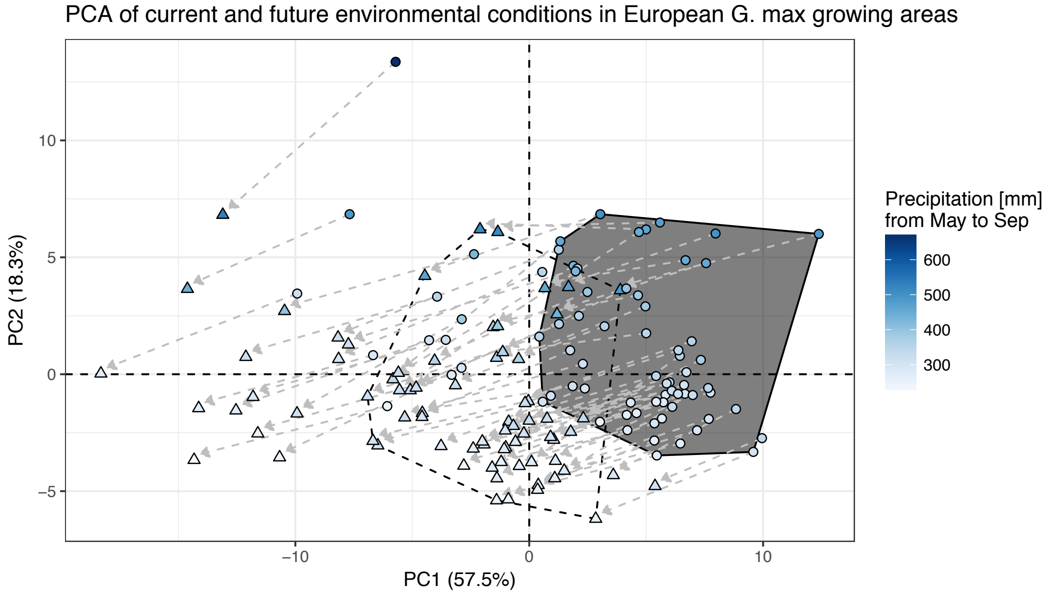


**Fig. S 8.** Distribution of Central (diamonds) and South (triangles) European soybean growing environments along the first two components of a PCA performed with environmental data characterizing the target population of environments. Colouring of sites was done according to the summed precipitation for the approximate soybean season from May to mid September. Arrows indicate the projected change of conditions until 2070. Scopes of Central European scenarios (today and in 2070) are indicated with polygons.

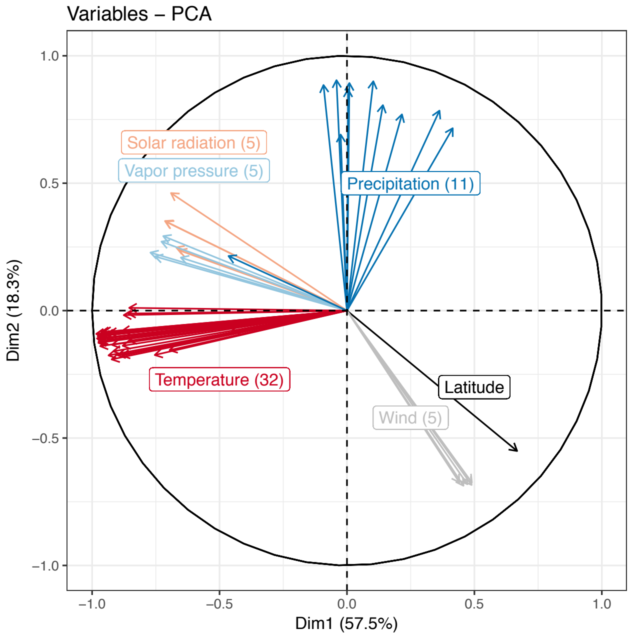


**Fig. S 9.** Correlation circle for the first two components of the PCA with environmental data characterizing European soybean growing environments. The correlation of a given variable with the first component can be read off from the x-axis and for the second component from the y-axis, respectively.

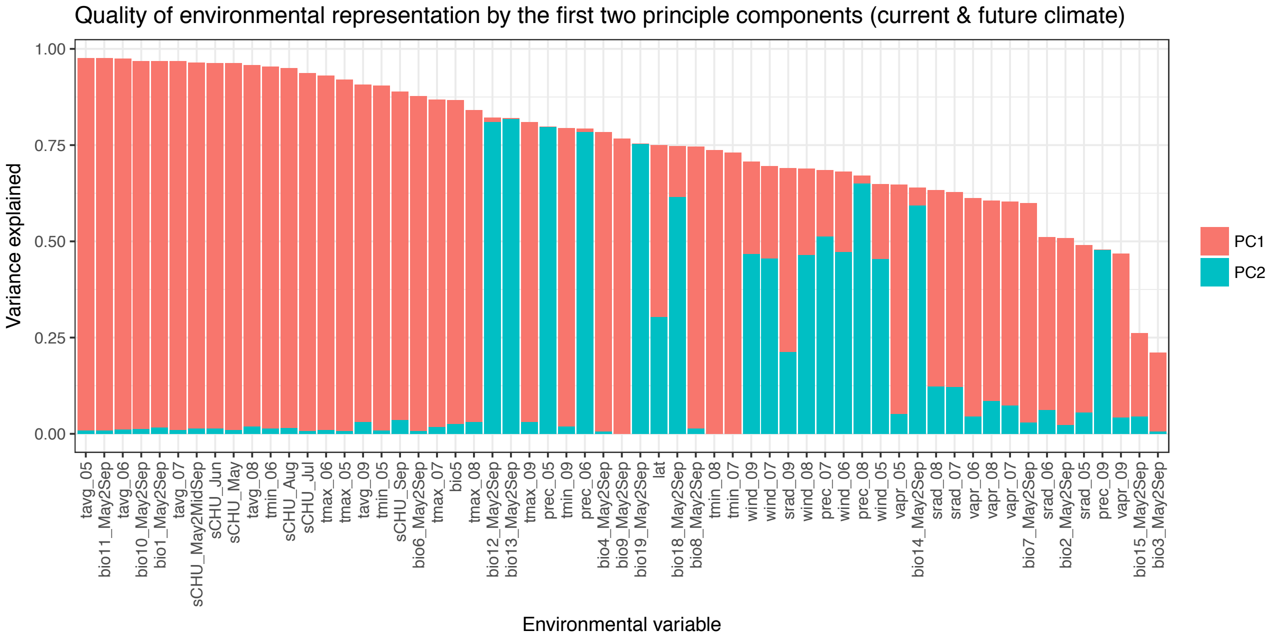


**Fig. S 10.** Quality of representation of original environmental variables characterizing European soybean growing environments jointly by the first two principal components.

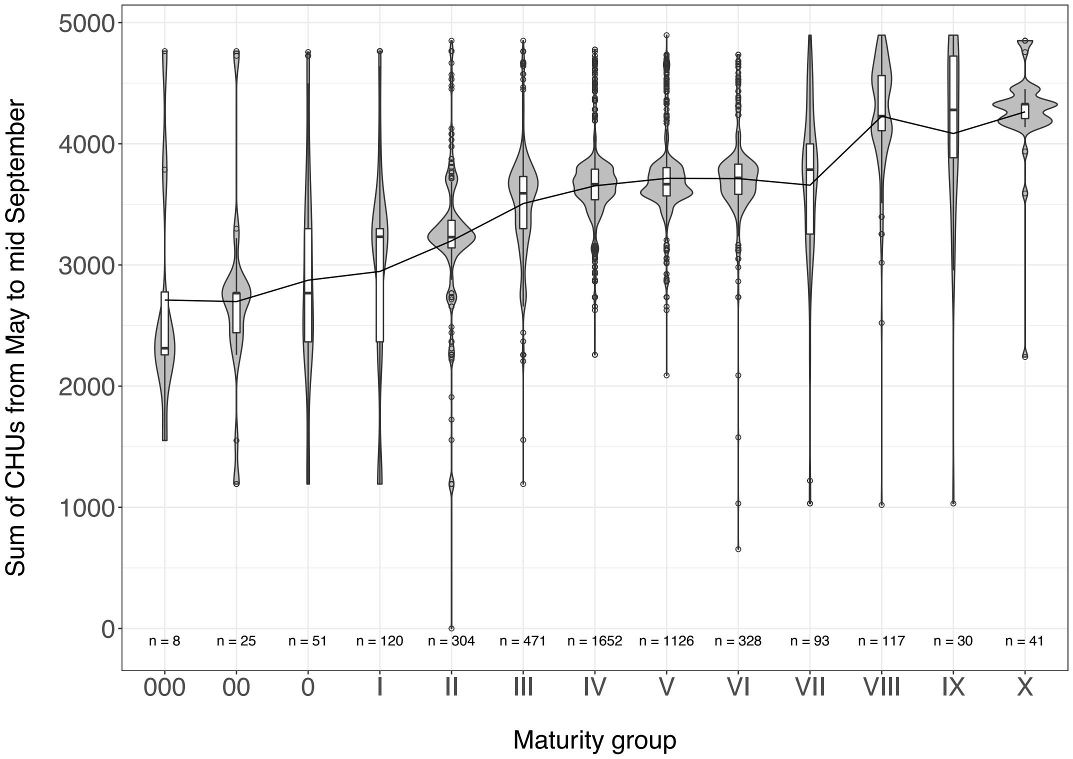


**Fig. S 11.** Violin / box plots of Asian georeferenced *G. max* accessions of different maturity groups and the respective sum of Crop Heat Units from May to mid September for the respective collection sites. The black line indicates the mean sum of CHUs within MGs and "n" gives the number of accessions for which MG and georeference / environmental data was available.

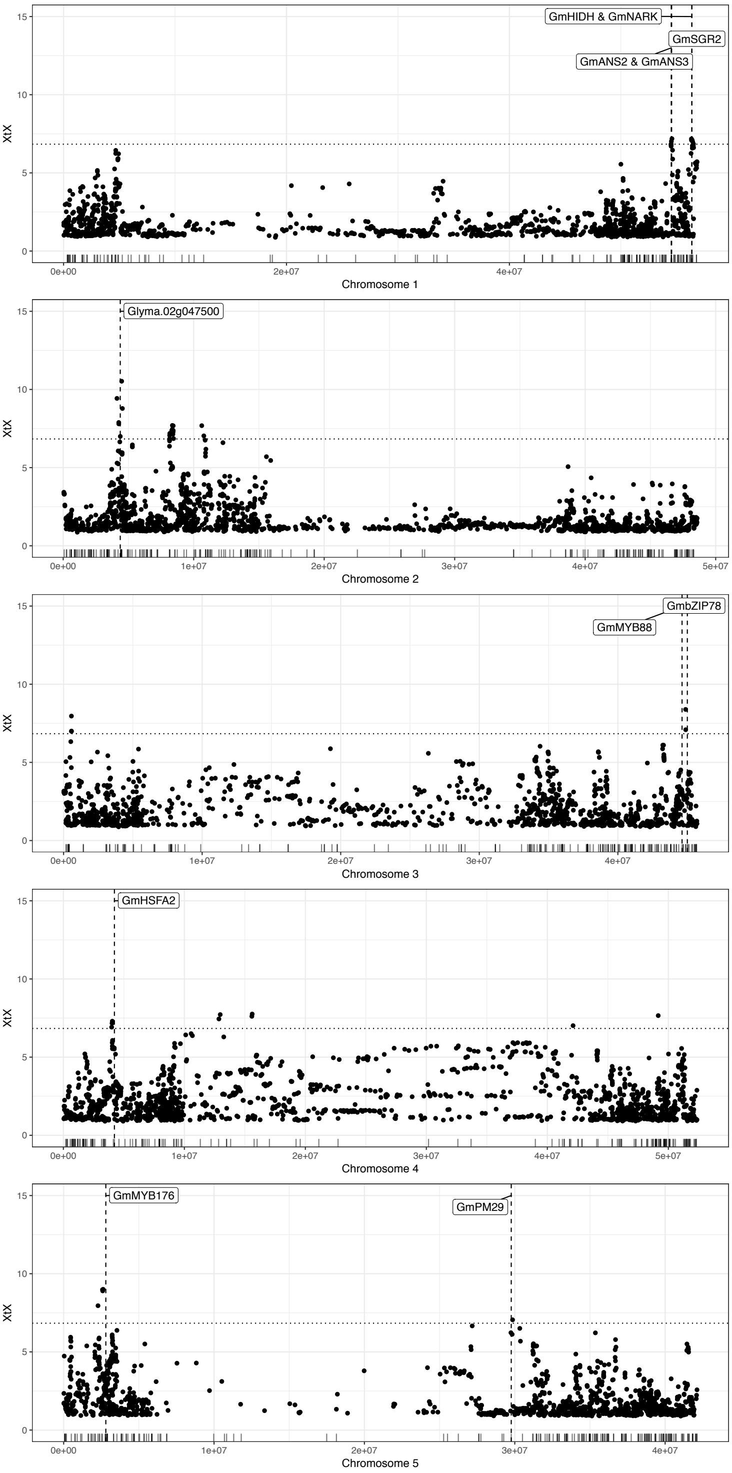


**Fig. S 12.** Pattern of differentiation between precore and non-candidate accessions along chromosomes as estimated with BAYPASS. Labels indicate genes, gene models and transcription factors that are related to abiotic adaptation and which were located within significantly differentiated regions (dotted horizontal line indicates the POD 99% quantile of *XtX* values). Rug indicates positions of published genes as deposited in www.soybase.org.

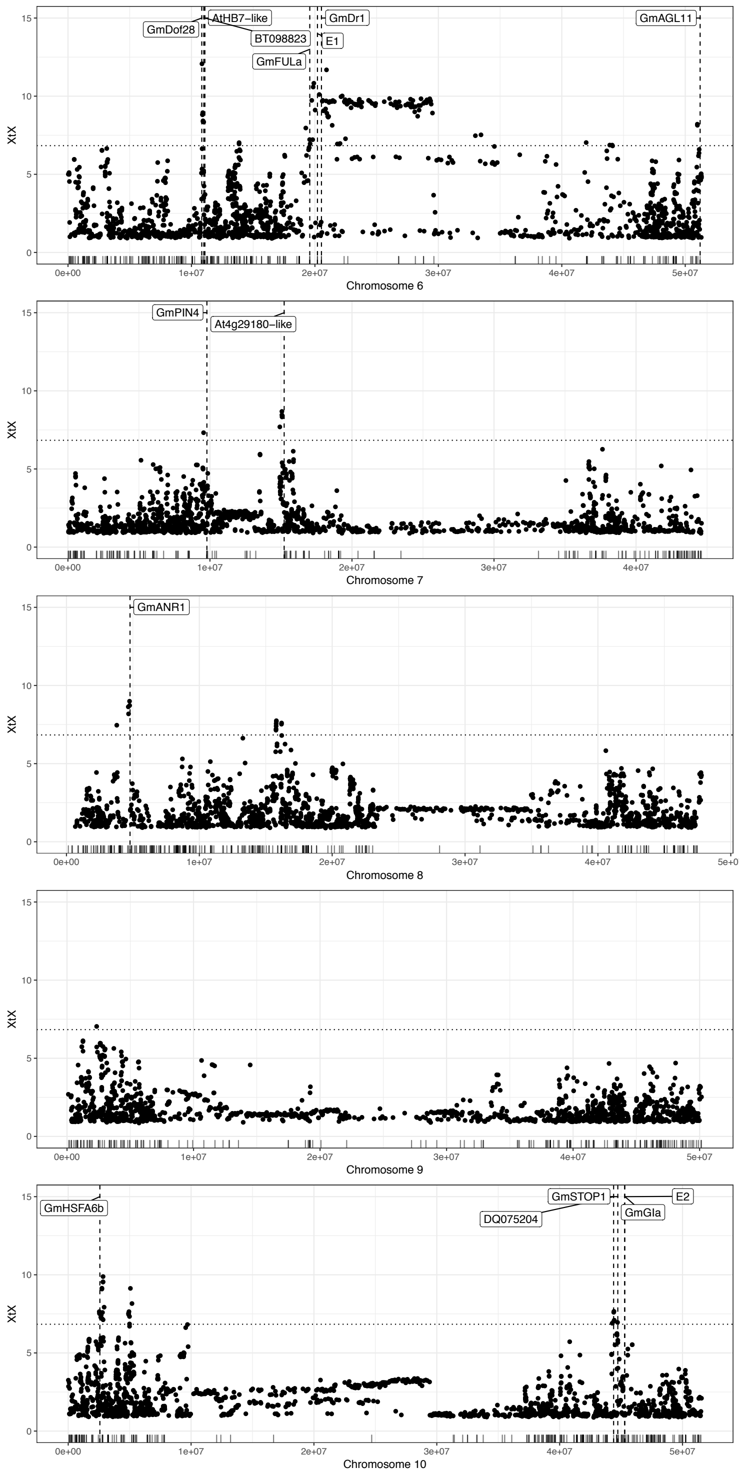


**Fig. S 13.** Pattern of differentiation between precore and non-candidate accessions along chromosomes as estimated with BAYPASS. Labels indicate genes, gene models and transcription factors that are related to abiotic adaptation and which were located within significantly differentiated regions (dotted horizontal line indicates the POD 99% quantile of *XtX* values). Rug indicates positions of published genes as deposited in www.soybase.org.

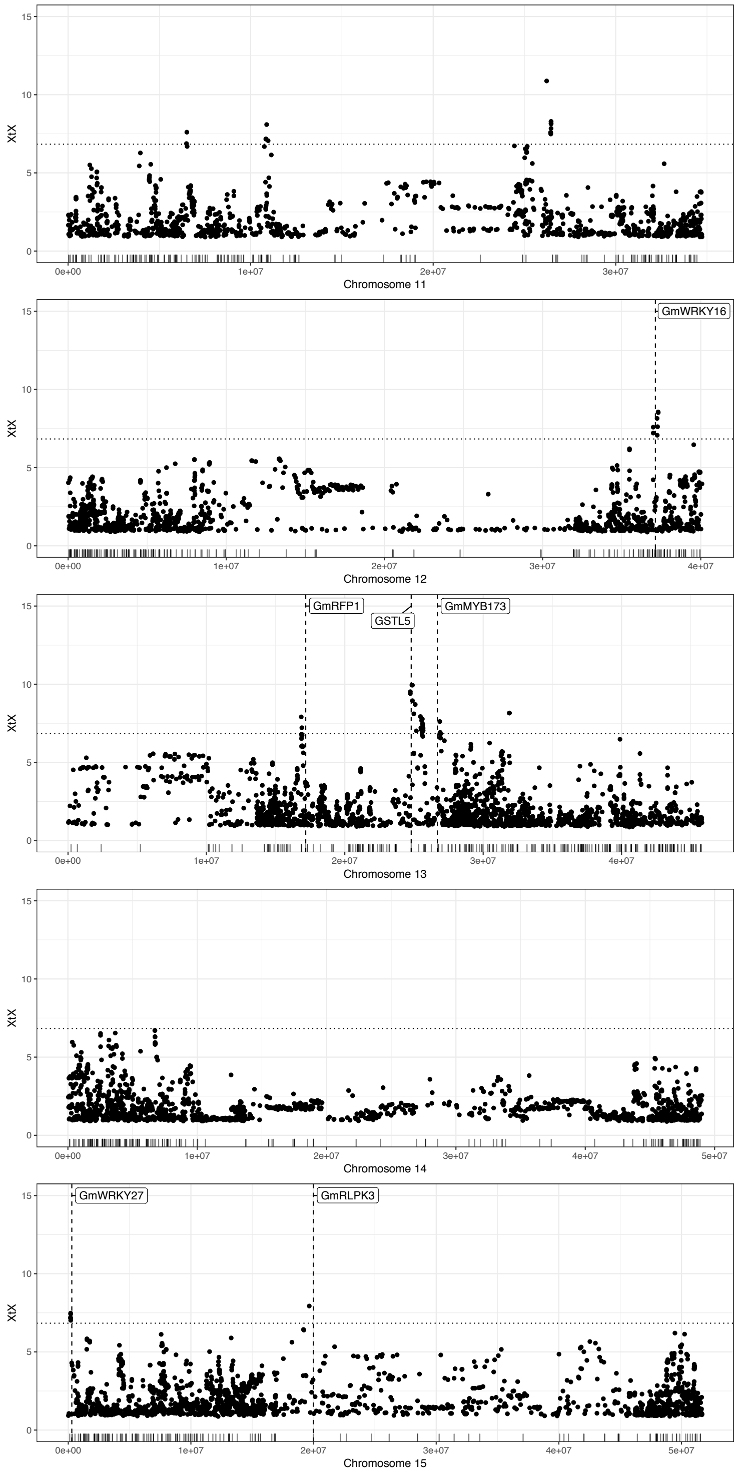


**Fig. S 14.** Pattern of differentiation between precore and non-candidate accessions along chromosomes as estimated with BAYPASS. Labels indicate genes, gene models and transcription factors that are related to abiotic adaptation and which were located within significantly differentiated regions (dotted horizontal line indicates the POD 99% quantile of *XtX* values). Rug indicates positions of published genes as deposited in www.soybase.org.

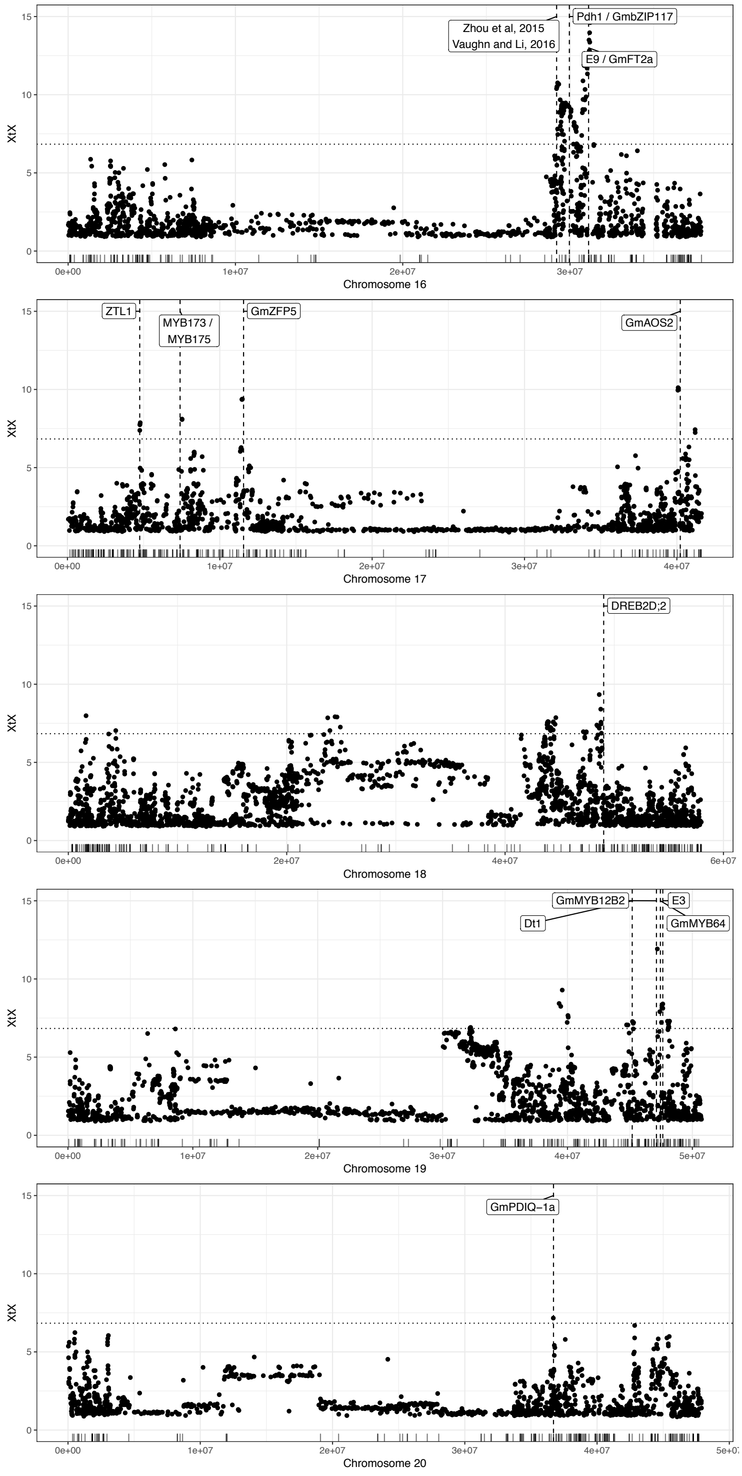


**Fig. S 15.** Pattern of differentiation between precore and non-candidate accessions along chromosomes as estimated with BAYPASS. Labels indicate genes, gene models and transcription factors that are related to abiotic adaptation and which were located within significantly differentiated regions (dotted horizontal line indicates the POD 99% quantile of *XtX* values). Rug indicates positions of published genes as deposited in www.soybase.org.

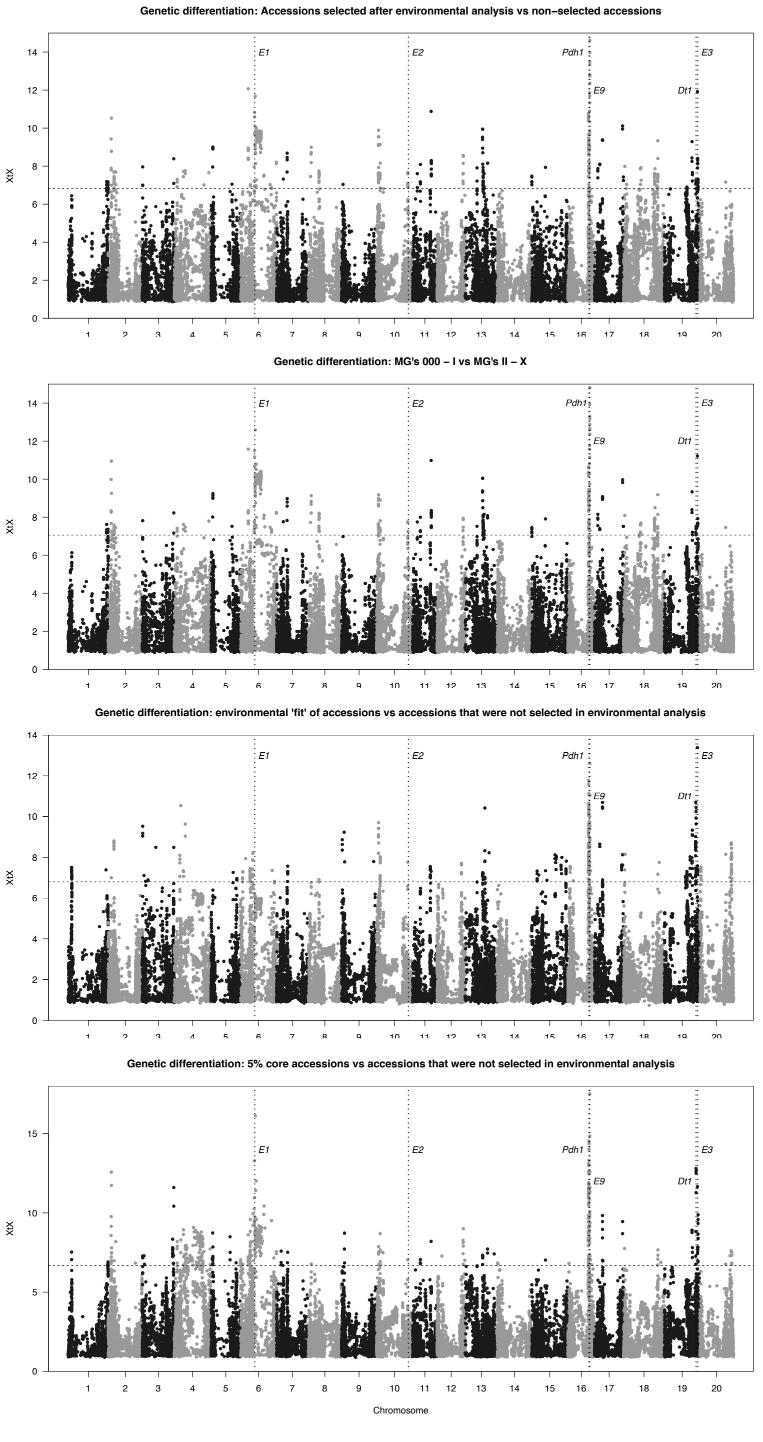


**Fig. S 16.** From top to bottom: Pattern of genome wide differentiation between precore and non-candidate accessions, early maturity accessions and late maturity accessions, precore accessions solely selected based on environmental data and accessions not selected based on environmental data (leaving out all accessions without eligible georeference information, no consideration of maturity group ratings) and 5%-core entries vs non-candidate accessions as estimated with BAYPASS. Dotted horizontal lines indicate the POD 99% quantile of *XtX* values. Positions of benchmark loci for environmental adaptation are indicated by vertical lines (*Dt1*, *E1*-*E3*, *E9* and *Pdh1*).

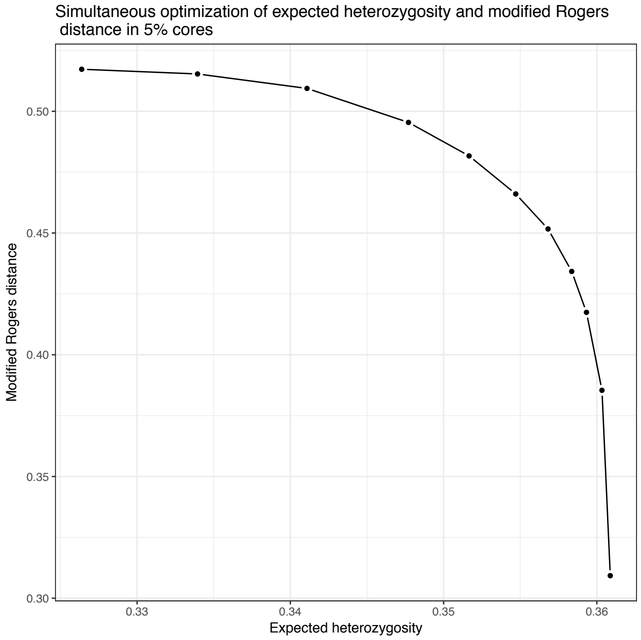


**Fig. S 17.** Pareto frontier of simultaneous optimization of allelic diversity and genetic distance in 5% cores with varying weights (0 to 1 in steps of 0.1). Allelic diversity was assessed by the expected heterozygosity and genetic diversity as the average of the minimum modified Rogers distance between accessions in cores.

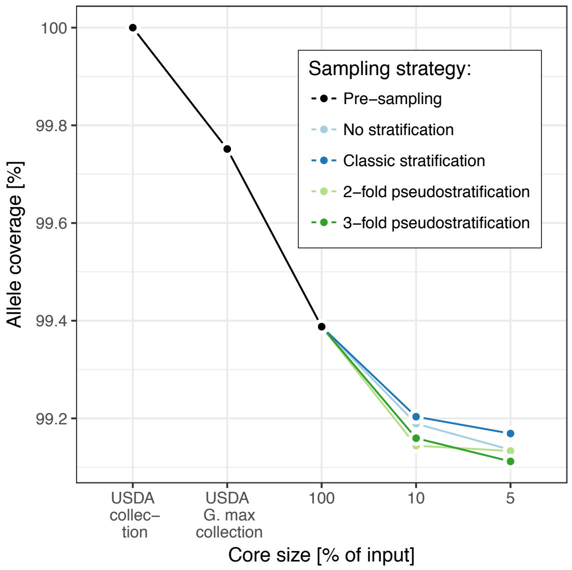


**Fig. S 18.** Decrease of allele coverage as a result of (1) exluding *G. soja* germplasm, (2) environmental stratification and selection of the precore and (3) the core collection formation.

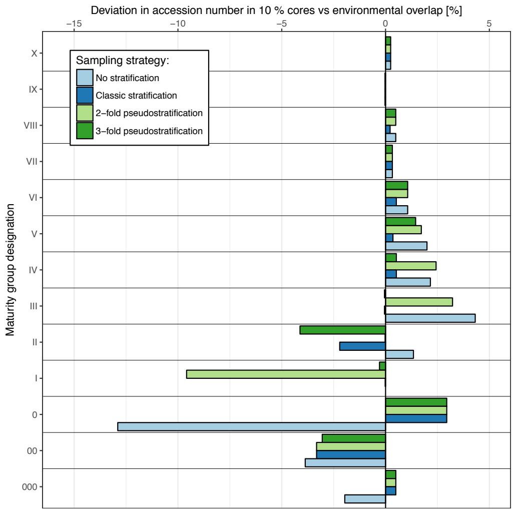


**Fig. S 19.** Evaluation of different core sampling strategies with regards to the preservation of MG fractions in 10% cores relative to the precore.

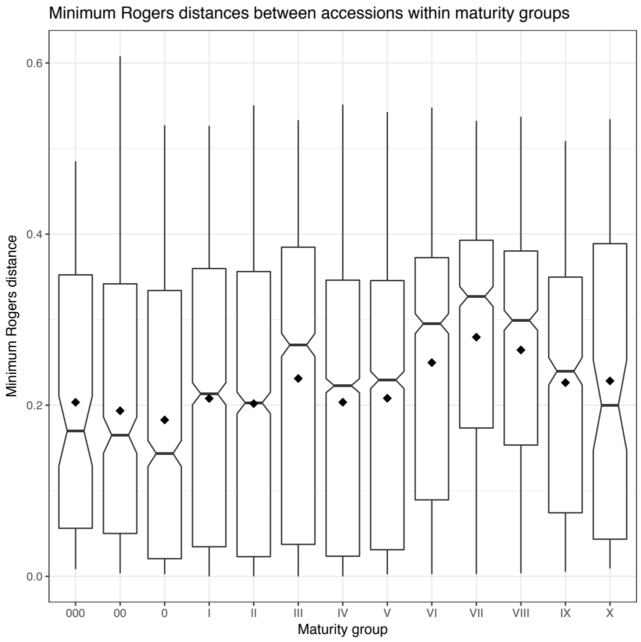


**Fig. S 20.** Minimum Rogers distances between accessions within maturity groups: Early maturity accessions on average were less distant from each other and thus were more prone to be eliminated from core solutions seeking to maximize this parameter (without stratification).

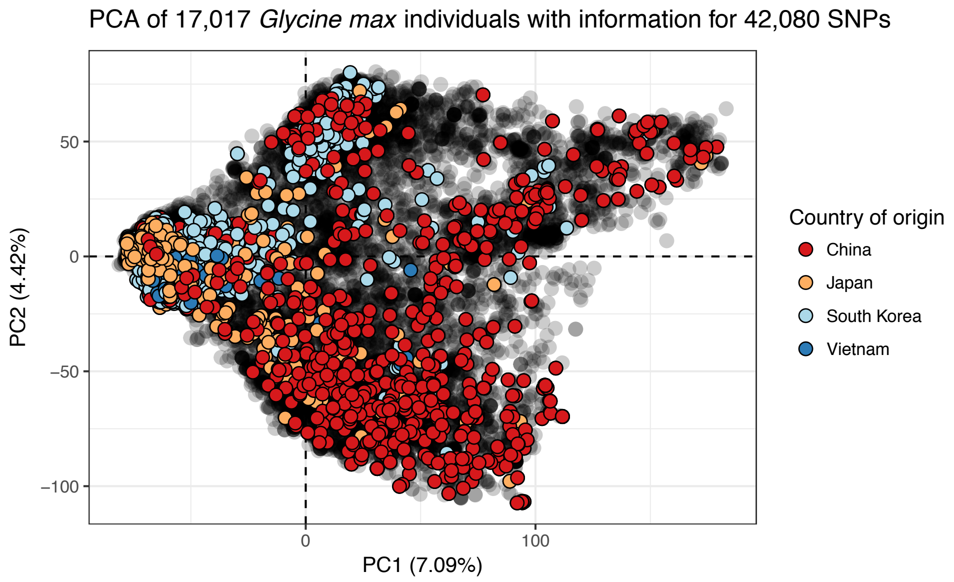


**Fig. S 21.** Principal component analysis summarizing the genetic structure in > 17.000 *G. max* accessions conserved in the *Introduced G. max* sub-collection of the USDA Soybean Germplasm Collection. Each dot represents one accession, colouring according to main provenances of germplasm (if recorded).

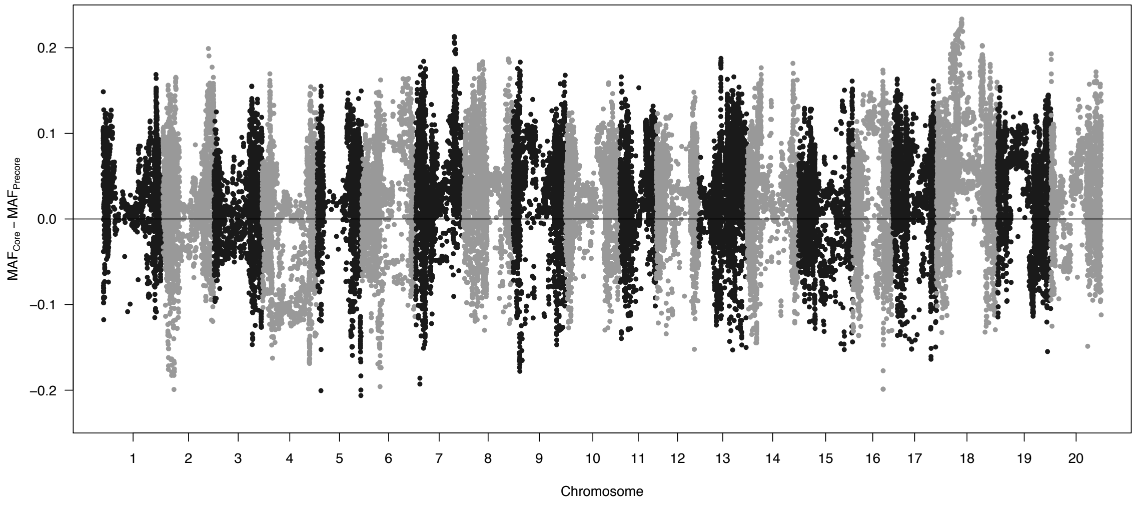


**Fig. S 22.** Genome wide changes in minor allele frequencies (MAFs): Each dot represents the MAF deviation in the 5% core compared to the precore at the respective marker. Dots above the horizontal line indicate SNP-wise elevations of MAF over the precore, dots below the digonal indicate decreases of MAF, respectively.
